# Evolutionary Dynamics of the Complete Chemosensory Repertoire in Kissing Bugs of the Genus *Rhodnius*: Divergent Odorant Receptors Contrast with Conserved Gene Families

**DOI:** 10.64898/2026.07.09.737527

**Authors:** Marie Merle, Gabin Rignault, Florence Mougel, Louise Maille, Jonathan Filée, Elaine Folly-Ramos, Carlos Eduardo Almeida, Myriam Harry

## Abstract

Chemosensory systems play a central role in host detection, feeding behavior, and habitat selection in hematophagous insects. Here, we performed a comparative evolutionary analysis of chemosensory gene repertoires across 13 species of the Chagas disease vector genus *Rhodnius*.

While gustatory receptors (GRs), ionotropic receptors (IRs), odorant-binding proteins (OBPs), and chemosensory proteins (CSPs) remained globally conserved, odorant receptors (ORs) displayed extensive lineage-specific expansions, tandem duplications, dynamic transcriptomic regulation, and recurrent signatures of positive selection. Major OR expansions were observed in *Rhodnius robustus* and *Rhodnius colombiensis*, suggesting increased sensory diversification in ecologically heterogeneous lineages. In contrast, conserved GR1 expression supports the maintenance of ancestral sugar-detection pathways despite hematophagy lifestyle. We further found no evidence of the canonical insect CO₂-associated GRs, suggesting alternative molecular mechanisms for CO₂ perception in Triatominae. Several receptors, including Orco, also displayed shifts in selective constraints between sylvatic and domiciliary species, consistent with sensory remodeling associated with adaptation to domestic habitats.

Together, our results identify ORs as the most evolutionarily dynamic component of the *Rhodnius* chemosensory repertoire and highlight contrasting evolutionary trajectories among chemosensory gene families during ecological diversification and vector adaptation.

## Introduction

Chemosensory systems are among the fastest evolving sensory pathways in insects and play a central role in ecological adaptation (Benton et al. 2006). Chemosensory genes belong to large multigene families whose repertoires frequently reflect host use, habitat and ecological specialization. In insects, odorant receptors (ORs), gustatory receptors (GRs), and ionotropic receptors (IRs) constitute the three main receptor families involved in odor and taste perception, while odorant-binding proteins (OBPs) and chemosensory proteins (CSPs) participate in ligand transport within sensilla (Hansson and Stensmyr 2011; Mitchell et al. 2020). Variation in the size and composition of these repertoires is primarily driven by gene duplication, pseudogenization, and functional divergence, resulting in major differences among species occupying distinct ecological niches.

The species of the genus *Rhodnius* (Hemiptera, Reduviidae, Triatominae) provide a relevant system to investigate the evolution of chemosensory gene families in relation to ecological diversification. These hematophagous bugs are vectors of *Trypanosoma cruzi*, the causative agent of Chagas disease. The genus currently includes 24 species, including the former *Psammolestes* species now nested within *Rhodnius* based on phylogenomic analyses (Filée et al. 2022). Most *Rhodnius* species are sylvatic and associated with palm trees, although some species also exploit bromeliads or other habitats (Abad-Franch et al. 2005), and feed on a broad range of vertebrate hosts (Ribeiro et al. 2014). Due to anthropogenic pressure disturbing or destroying natural habitats, some species have colonized human dwellings and transmit the parasite to humans via the infected feces.

Comparative studies across insects have linked chemosensory evolution to ecological specialization. Generalist species exposed to chemically diverse environments usually possess larger repertoires than specialists, whereas ecological shifts are frequently associated with lineage-specific expansions, gene losses, or signatures of selection in chemosensory receptors. For example, the specialist *Drosophila sechellia* exhibits accelerated loss of OR and GR genes relative to its generalist sister species *D. simulans* (McBride 2007), while odorant and gustatory receptor genes show strong differentiation among host races of the pea aphid *Acyrthosiphon pisum* (Smadja et al. 2012).

Hematophagy is expected to impose strong constraints on chemosensory evolution. Chemosensory repertoires have been partially characterized in hematophagous Hemiptera. In the domiciliary species *R. prolixus*, 116 ORs (including 5 annotated as pseudogenes) and 28 GRs were identified from the genome (Mesquita et al. 2015), whereas the highly specialized bed bug *Cimex lectularius* possesses approximately 50 ORs and 35 GRs (Hansen et al. 2014). In *R. prolixus*, lineage-specific expansions of several chemosensory families were proposed to be associated with hematophagy (Mesquita et al. 2015). Functional studies further support the ecological relevance of receptor diversification in this genus, heterologous expression assays identified ligand specificity for an OR (Franco et al. 2018). Contrastingly, a GR named RproGR1 appears conserved with fructose receptors such as DmGR43a in *Drosophila melanogaster* and AmGR3 in *Apis mellifera* (Mesquita et al. 2015). Interestingly, *R. prolixus* can feed on plant substrates under laboratory conditions (Díaz-Albiter et al. 2016), and palm DNA was recently detected in the gut contents of *R. robustus* (Da Lage et al. 2024), suggesting retention of ancestral sugar-related functions and questioning the obligate hematophagy of Triatominae. Changes in gene expression may also contribute to ecological adaptation. In Triatominae, chemosensory genes are underexpressed in domiciliary populations relative to sylvatic populations of *Triatoma brasiliensis* (Marchant et al. 2016), and several receptors are specifically expressed in the sylvatic species *R. robustus* compared with the domiciliary species *R. prolixus* (Marchant et al. 2021). These observations suggest that both repertoire evolution and regulatory divergence contribute to adaptation to distinct ecological environments.

Despite these advances, a comparative evolutionary framework integrating multiple *Rhodnius* species is still lacking. It remains unclear which components of the chemosensory repertoire remained conserved across the genus and which evolved through lineage-specific diversification in relation to ecological specialization. To fill this gap, using genomic data from 13 *Rhodnius* species and transcriptomic data for 9 of them, we combined comparative annotation, orthology reconstruction, phylogenetic analyses, and tests of selection to investigate the evolution of major chemosensory gene families in relation to ecological diversification.

## Materials and Methods

### Taxon Sampling and Genome Data

Genome assemblies for all species were previously generated and described in Merle et al. (submitted) and were retrieved for downstream analyses (Table 1).

**Table 1.** Summary of genomic and transcriptomic resources: genome assembly metrics (completeness, length, contig count), and DOI or accession numbers for each dataset.

| Species | Sample name | Genome assembly | Busco S (%) | Total ungapped length (kb) | Contigs (#) | RNA-seq |
| --- | --- | --- | --- | --- | --- | --- |
| *R. brethesi* | RBAM | [zenodo.18352118](https://zenodo.org/record/18352118) | 88.4 | 468,07 | 10,466 | [SRR36983270](https://srr36983270) |
| *R. colombiensis* | RCOLM | [zenodo.18352118](https://zenodo.org/record/18352118) | 95.1 | 479,825 | 7,465 | [SRR36983275](https://srr36983275) |
| *R. coreodes* | PSAMc | [JBTXIK000000000](https://doi.org/10.5281/zenodo.18352118) | 95 | 579,397 | 5,000 | - |
| *R. domesticus* | RDOMFM | [zenodo.18352118](https://zenodo.org/record/18352118) | 94.7 | 538,754 | 3,910 | - |
| *R. milesi* | RMM | [zenodo.18352118](https://zenodo.org/record/18352118) | 93.1 | 522,085 | 4,008 | - |
| *R. montenegrensis* | R8F | [zenodo.18352118](https://zenodo.org/record/18352118) | 79.6 | 353,110 | 10,354 | [SRR36983273](https://srr36983273) |
| *R. nasutus* | R7WU | [zenodo.18352118](https://zenodo.org/record/18352118) | 61.4 | 269,815 | 11,363 | [SRR36983269](https://srr36983269) |
| *R. neglectus* | R5WU | [zenodo.18352118](https://zenodo.org/record/18352118) | 86.6 | 420,276 | 9,263 | [SRR36983274](https://srr36983274) |
| *R. neivai* | RNEIF | [zenodo.18352118](https://zenodo.org/record/18352118) | 91.5 | 676,140 | 17,255 | - |
| *R. pallescens* | RPALM | [zenodo.18352118](https://zenodo.org/record/18352118) | 91.2 | 410,473 | 11,565 | [SRR36983272](https://srr36983272) |
| *R. pictipes* | RPICM | [zenodo.18352118](https://zenodo.org/record/18352118) | 86 | 462,908 | 11,624 | [SRR36983271](https://sra.nlm.nih.gov/submit/srr/SRR36983271) |
| *R. prolixus* | ProH2 | [JBTXIJ0000000000](https://jbt.xij0000000000) | 88 | 474,000 | 13,347 | [ERR1315264](https://sra.nlm.nih.gov/submit/err/ERR1315264) |
| *R. robustus* | RRTM | [JBTXII0000000000](https://jbt.xii0000000000) | 89.8 | 570,799 | 6,344 | [ERR1315270](https://sra.nlm.nih.gov/submit/err/ERR1315270) |

### Chemosensory Gene Annotation

Annotation was performed using previously described chemosensory genes (GRs, ORs, IRs, OBPs, and CSPs) from hemipteran species, including those annotated in the *R. prolixus* genome (see Supplementary Table 1 for detailed references of query sequences for each gene family).

Genome-wide searches were conducted using the protein2genome model implemented in Exonerate (v 2.4.0; Slater and Birney 2005) with the maximum intron length parameter set to 3,000 bp. Given the birth-and-death evolution of chemosensory receptor genes and their occurrence in tandem arrays, predicted OR, GR, and IR candidates were further processed using InsectOR (Karpe, Tiwari, and Ramanathan 2021) with default parameters (alignment cluster cutoff = 1; completion cutoff = 300 amino-acids). Due to the homology between glutamate receptors and IRs, we carefully filtered out glutamate receptor hits from IR annotations using BLAST against the non-redundant database and phylogenetic inference. Note that InsectOR does not identify alternative transcripts, which were therefore not considered in this analysis. Using the presence of frameshifts or stop codons and the cutoff used, each sequence is considered as normal, pseudogene, complete, or partial by InsectOR.

OBPs and CSPs were annotated through similarity searches against genome assemblies using tblastn (v. 2.10; Altschul et al. 1990) and Exonerate with the protein2genome model (v. 2.4.0). Annotated genes from *R. prolixus* and other hemipteran species were used as queries, and significant matches were retained using an e-value threshold < 1e-5 and a score > 300.

### Gene Phylogenies

Amino acid sequence alignments were performed for each gene family dataset using MAFFT (Katoh, Rozewicki, and Yamada 2019). Maximum-likelihood trees were inferred using IQ-TREE (Nguyen et al. 2015) with 1,000 ultrafast bootstrap replicates (Minh, Nguyen, and Von Haeseler 2013). Substitution models for each gene family were selected using ModelFinder (Kalyaanamoorthy et al. 2017) and are reported in the corresponding tree legends.

Due to the absence of CO_2_ receptors in the GR repertoire (Mesquita et al. 2015), which are typically used for rooting, the GR tree was rooted using the midpoint method. This method was also used for CSP and OBP trees. The OR and IR phylogenies were rooted using ORco and IR25a, respectively. Trees were visualized using iTOL (Letunic and Bork 2019).

### Molecular Evolution Analysis

Selective pressures were estimated using the dN/dS ratio (ω) in CodeML (PAML, Yang 2007) through EasyCodeML (Gao et al. 2019) to perform likelihood ratio tests (LRTs) between nested models.

Orthologous groups were defined from phylogenies based on well-supported groups (single-copy genes per species, or up to two copies in one species). Groups with poor alignments (e.g., extensive gaps or ambiguous regions), or species-specific expansions were excluded. Duplicates were analyzed separately to avoid conflating paralog-specific evolutionary signals.

Positive selection was assessed using M1a vs M2a, M7 vs M8, and M8a vs M8, where alternative models allow a class of codons with ω > 1. A branch model tested differences in ω between habitat categories (domiciliary vs sylvatic; Filée et al. 2022) with branches assigned accordingly, by comparing a two-ratio to a one-ratio null model. Site-specific variation in selective pressure was also tested using M0 vs M3.

P-values were corrected for multiple testing using the Benjamini-Hochberg procedure (Benjamini and Hochberg 1995), with adjusted P < 0.05 considered significant.

### RNA-seq Quantification

Transcriptomic data from seven *Rhodnius* species were analyzed to quantify gene expression. RNA-seq datasets from head, rostrum, and antennal tissues were sourced from Marchant et al. (2021) and Merle et al. (submitted) (Table 1). A unified reference index was built to ensure inter-specific comparison. This index comprised the coding sequences (CDS) of *R. prolixus* (Latorre-Estivalis et al. 2017), supplemented with new sequences identified in this study (labeled with the suffix “_NEW”) or with the closest orthologs when absent. Transcript abundance was quantified using Kallisto v0.46.2 (Bray et al. 2016) and normalized as Transcripts Per Million (TPM). For genes showing differential dN/dS ratio, a close-up heatmap comprising only these genes was performed, normalized by the mean of expression across all the species, ensuring the visualization.

## Results

### Chemosensory Gene Annotation

To account for differences in assembly quality and annotation completeness among species, chemosensory gene repertoires were analyzed by distinguishing complete genes (with start codon), partial genes, and pseudogenes. The “total number” of genes refers to the combined set of these three categories. Comparative analyses between species were primarily based on complete genes (Table 2), whereas partial genes and pseudogenes were considered as complementary indicators of repertoire diversity and turnover (Supplementary Table 2). All annotated genes are expressed in the head of at least one *Rhodnius* species, confirming the annotation (Supplementary Figures 1-5).

**Table 2:**
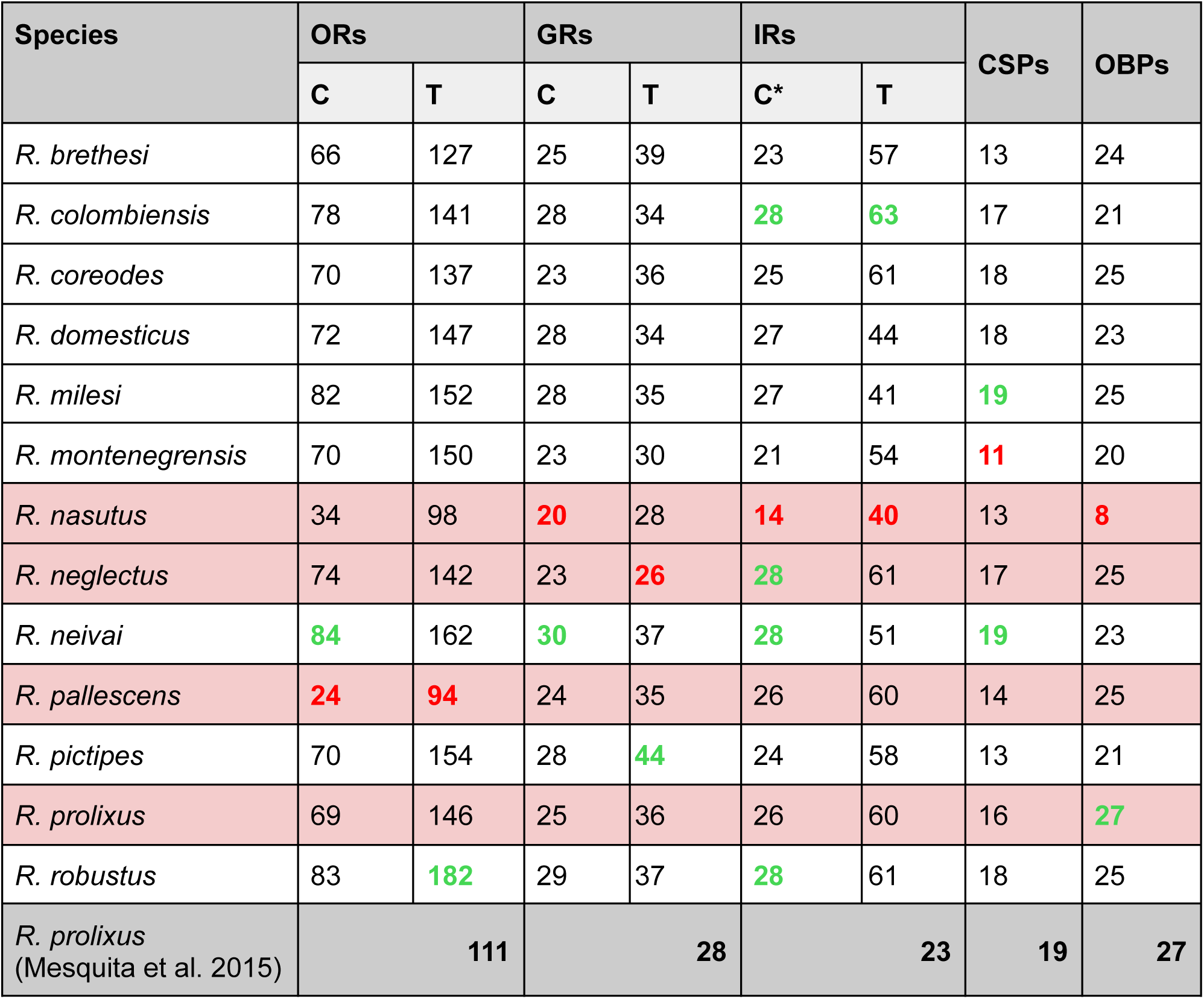
Results of chemosensory gene annotation. Domiciliary species are colored in red. C: Complete, non-pseudogenized, with start codon; T: Includes partial genes and pseudogenes. REF: Reference data from Mesquita et al. (2015). *: After manual curation. Highest values are written in bold and red and lowest in bold and green.

The total number of ORs identified in our assemblies varied substantially among species, ranging from 94 in *R. pallescens* to 182 in *R. robustus*. The proportion of complete genes containing a start codon was also highly variable, from 24 in *R. pallescens* to 84 in *R. neivai*, whereas partial sequences represented a substantial fraction of the repertoire in several species. Pseudogene content remained limited, ranging from 3 in *R. pallescens* to 22 in *R. coreodes* (Supplementary Table 2). Among domiciliary species, complete OR repertoires ranged from 24 to 74 genes, *R. nasutus* and *R. pallescens* exhibit the smallest complete OR repertoires (<35 ORs), whereas sylvatic species ranged from 66 to 84 OR genes, with *R. neivai* and *R. robustus* displaying the largest OR repertoires (>80 ORs). Duplication events were observed in 13 ORs, of which 9 were restricted to sylvatic species (Supplementary Table 3).

For the GRs identified, the total number of genes varied moderately among species, ranging from 26 in *R. neglectus* to 44 in *R. pictipes*. The number of complete GRs (with a start codon) ranged from 20 *in R. nasutus* to 30 in *R. neivai*, whereas partial sequences represented a variable fraction of the repertoire depending on the species. The reduced genome completeness of *R. nasutus* may partly explain its lower GR count. Pseudogene content remained limited, ranging from 0 in *R. nasutus* and *R. neglectus* to 5 in *R. prolixus* and *R. coreodes* (Supplementary Table 2). Among domiciliary species, complete GR repertoires ranged from 20 to 25 genes, whereas sylvatic species ranged from 23 to 30 GR genes. Notably, among sylvatic species, *R. neivai* and *R. robustus* displayed the largest GR repertoires, while *R. coreodes* and *R. montenegrensis* displayed the smallest ones.

For the IRs, the total number of genes ranged from 40 in *R. nasutus* to 63 in *R. colombiensis*, whereas the number of complete IRs containing a start codon and manually checked ranged from 14 in *R. nasutus* to 28 in *R. colombiensis*, *R. neglectus*, *R. neivai* and *R. robustus*. As for GR genes, the reduced genome completeness of *R. nasutus* may partly explain its lower IR count. Partial sequences ranged from 4 in *R. neivai* to 23 in *R. prolixus* and pseudogene content from one in *R. montenegrensis* and *R. pictipes* to 10 in *R. coreodes* and *R. robustus* (Supplementary Table 2). Excluding *R. nasutus*, the domiciliary species exhibited IR repertoires ranging from 26 to 28 IR genes and the sylvatic species, from 21 to 28 genes.

For the annotation of additional OBP and CSP genes, we employed a different pipeline, which did not provide detailed classification of pseudogenes or partial sequences. Nevertheless, all annotated genes, which are smaller in size, were complete. The number of CSPs ranged from 11 in *R. montenegrensis* to 19 in *R. milesi* and *R. neivai*. The number of OBPs ranged from 20 in *R. montenegrensis* to 27 in *R. prolixus* Honduras, with the exclusion of *R. nasutus* due to the low quality of the genome. For reference, Mesquita et al. (2015) annotated 209 functional chemosensory genes in *R. prolixus*, comprising 111 ORs, 28 GRs, 26 IRs, 16 CSPs, and 27 OBPs. In our study including 13 *Rhodnius* species, we identified 11 new orthologous groups, including 7 OR orthogroups, 1 GR orthogroup, 2 IR orthogroups, and 1 CSP orthogroup (Supplementary Tables 3-7).

### Orthologous Groups and Phylogeny

Phylogenetic analyses were conducted using complete genes (with a start codon) and including all previously annotated *R. prolixus* sequences. For each chemosensory gene family, the phylogeny enabled the definition of ortholog groups and the numbering of genes according to the nomenclature established by Mesquita et al. (2015) and completed by Latorre-Estivalis et al. (2016). The absence of some genes from the assemblies should be interpreted with caution. Although such patterns may reflect evolutionary events, they may also result from technical limitations leading to incomplete genome annotations, due to low sequencing coverage, or reduced assembly completeness.

The OR phylogeny enabled the definition of 42 unambiguous ortholog groups containing at least one representative from each species (Figure 1; Supplementary Table 3). Several OR groups showed evidence of lineage-specific expansions, notably OR21-23, OR28-29, and OR54, characterized by tandem duplications typical of recent expansion events. The OR28-29 group was particularly expanded, with 49 genes identified across the 13 assemblies, whereas OR21-23 comprised 25 genes. Within the OR28-29 group, six copies in the sylvatic species *R. colombiensis* and seven in *R. robustus* are organized in tandem on the same contig. Similar tandem organizations involving three genes were also observed for OR21-23 and OR54, notably in *R. neglectus*, and *R. milesi*.

**Figure 1:**
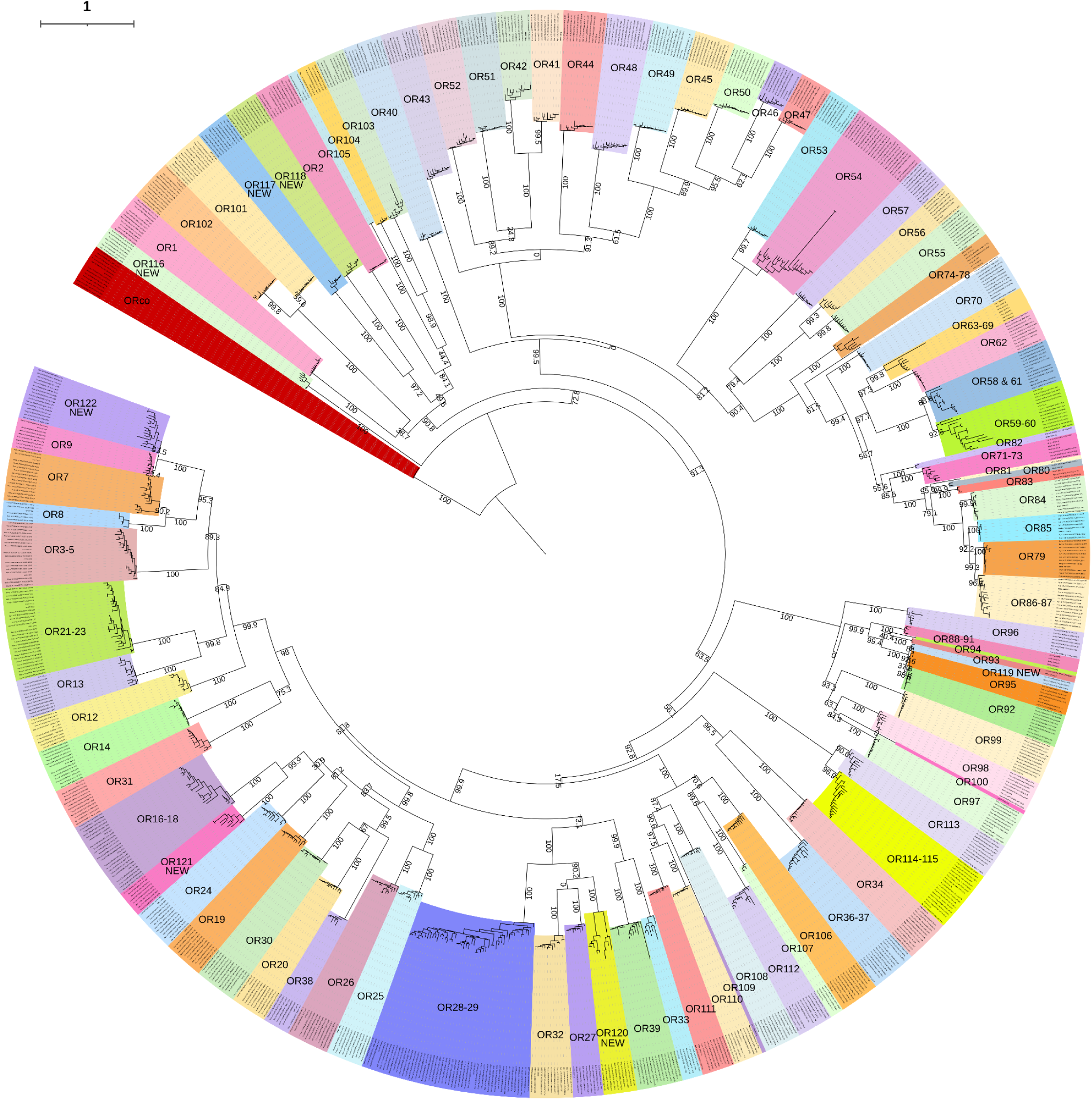
Consensus tree built from amino acid sequences of ORs from 13 *Rhodnius* species. The phylogeny was built with IQ-TREE using the maximum likelihood approach. The model used is JTT+F+G4. The bootstrap procedure was applied with 1,000 ultrafast bootstraps. ORs are numbered according to Mesquita et al. (2015). Bootstrap values are indicated on branches except within defined orthologous groups, where they have been removed for clarity. The species included are: reference *R. prolixus* (Mesquita et al. 2015), *R. brethesi*, *R. colombiensis*, *R. coreodes*, *R. domesticus*, *R. milesi*, *R. montenegrensis*, *R. nasutus*, *R. neglectus*, *R. neivai*, *R. pallescens*, *R. pictipes*, *R. prolixus* and *R. robustus*. The scale bar corresponds to 1 substitution per site.

Compared with the published *R. prolixus* repertoire, seven new ortholog groups (OR116 to OR122) were identified, whereas 18 previously described ORs were not recovered in the assemblies, namely OR63 to 69, OR81, OR75 to 78, OR88 to 91, OR94, OR100 and OR109. This discrepancy could be explained by several factors: the published sequences might correspond to pseudogenes or misannotated introns, or these regions might be missing due to gaps in the genome assembly, which remains highly fragmented. Conversely, several ORs described as partial in the *R. prolixus* reference were recovered as complete orthologs in other species, including OR71-73 in *R. neivai,* OR74 in *R. milesi* and *R. nasutus*, and OR85 in seven assemblies. These results highlight the benefit of comparative annotation approaches when InsectOR is used for improving gene reconstruction in multigene families.

The GR phylogeny revealed a much higher level of conservation, with most ortholog groups including one gene from each species and displaying short terminal branches (Figure 2; Supplementary Table 4). Moreover, the GR1 was found in all assemblies and showed a high expression in the nine *Rhodnius* species included in the expression heatmap (Supplementary Figure 2).

**Figure 2:**
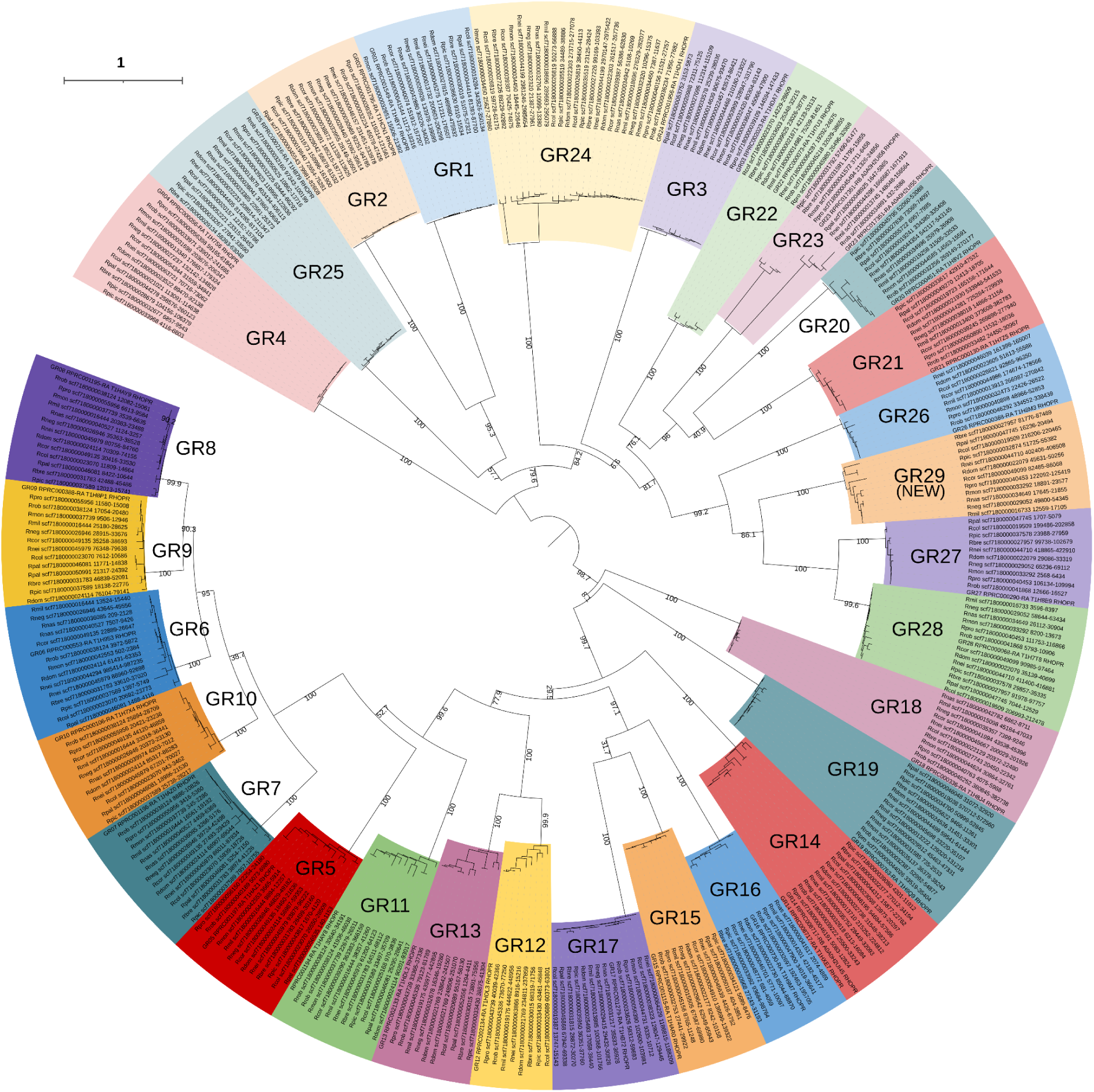
Consensus tree built from amino acid sequences of GR from 13 *Rhodnius* species. The phylogeny was built with IQ-TREE using the maximum likelihood approach. The model used is JTT+F+I+G4. The bootstrap procedure was applied with 1,000 ultrafast bootstraps. GRs are numbered according to Mesquita et al. (2015). Bootstrap values are indicated on branches except within defined orthologous groups, where they have been removed for clarity. The species included are: reference *R. prolixus* (Mesquita et al. 2015), *R. brethesi*, *R. colombiensis*, *R. coreodes*, *R. domesticus*, *R. milesi*, *R. montenegrensis*, *R. nasutus*, *R. neglectus*, *R. neivai*, *R. pallescens*, *R. pictipes*, *R. prolixus* and *R. robustus*. The scale bar corresponds to 1 substitution per site.

A novel ortholog group, GR29, absent from the original *R. prolixus* annotation, was identified in all other species and clustered with GR27 and GR28 in a tandem organization, suggesting recent paralogous duplications. In addition, multiple copies of GR24 were detected in 11 species, confirming a *Rhodnius*-specific expansion previously suggested by Mesquita et al. (2015).

A total of 34 ortholog groups were identified for IRs, most of which followed a conserved 1:1 orthology pattern (Figure 3; Supplementary Table 5). Unexpectedly, multiple duplications of the conserved co-receptor IR8a were detected, particularly in *R. colombiensis*, where up to six copies were identified. Because IR phylogenies are traditionally rooted using both IR8a and IR25a co-receptors, this atypical expansion prevented the use of IR8a as a stable reference, and the tree was therefore rooted using only the IR25a clade. Although assembly artifacts cannot be fully excluded, the recurrence of these duplications across several assemblies suggests an unusual evolutionary trajectory for IR8a within *Rhodnius*. In addition, major expansions were observed within the divergent IR75 clade (IR75a–IR75p), whereas IR107 appeared highly lineage-specific and was only recovered in *R. robustus* and the *R. prolixus* reference genome.

**Figure 3:**
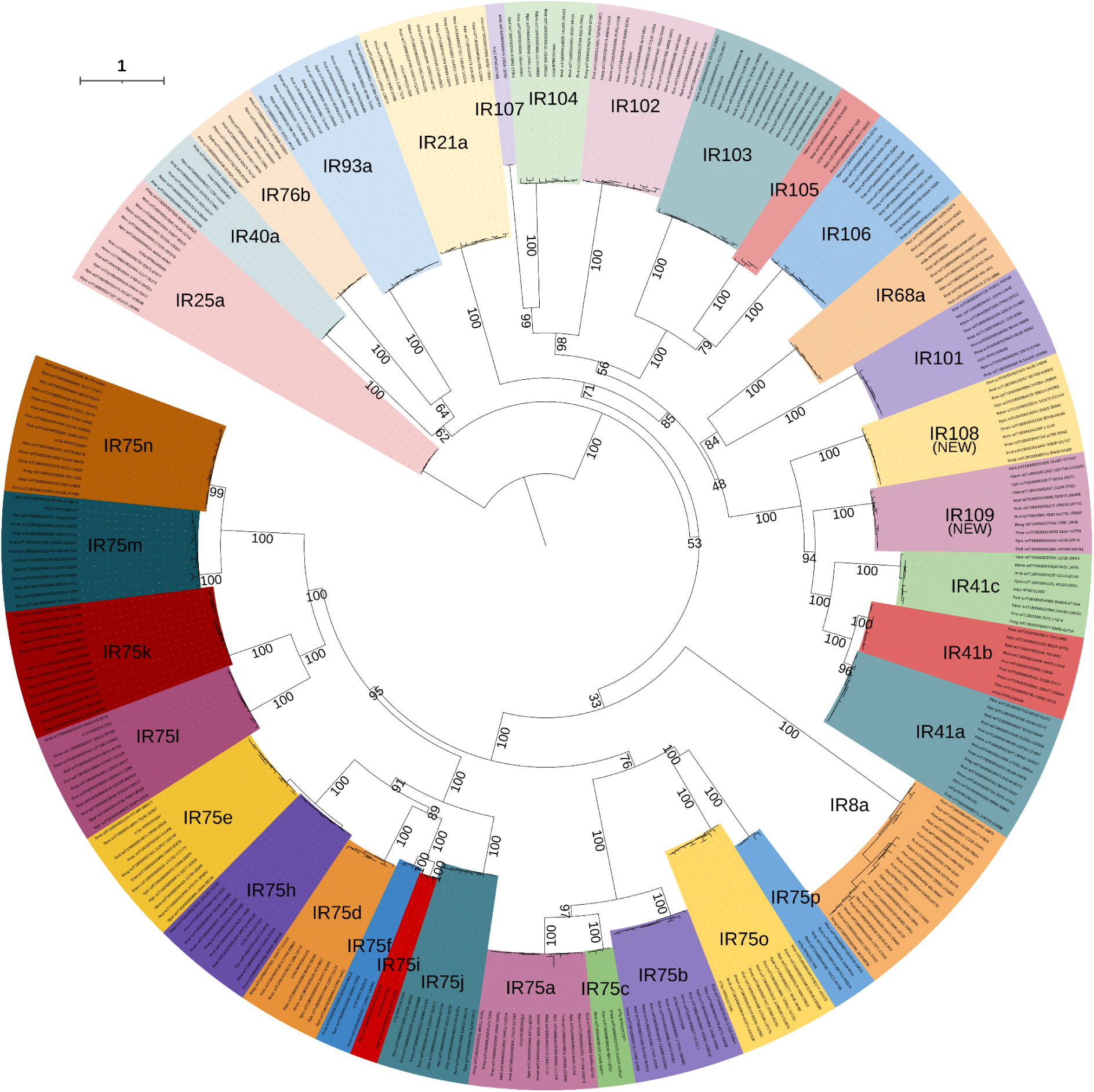
Consensus tree built from amino acid sequences of IR from 13 *Rhodnius* species. The phylogeny was built with IQ-TREE using the maximum likelihood approach. The model used is JTT+F+I+G4. The bootstrap procedure was applied with 1,000 ultrafast bootstraps. IRs are numbered according to Mesquita et al. (2015). Bootstrap values are indicated on branches except within defined orthologous groups, where they have been removed for clarity. The scale bar corresponds to 1 substitution per site.

Phylogenetic analyses of OBPs and CSPs confirmed a generally high level of conservation, with most ortholog groups including representatives from nearly all species (Figures 4 and 5; Supplementary Table 6 and 7). Twenty-seven OBP ortholog groups and 18 CSP ortholog groups were identified. A few OBP groups displayed more restricted distributions, and were limited to three species belonging to the *prolixus* group namely for OBP15 to *R. prolixus*, *R. robustus*, and *R. coreodes*, and for OBP3 to *R. prolixus*, *R. montenegrensis*, and *R. robustus,* suggesting recent lineage-specific expansions. CSP14-15 exhibited a complex history, with four copies identified in *R. prolixus* and contrasting expression profiles between paralogs, with one CSP14 lowly expressed and one CSP15 highly expressed (Supplementary Figure 5). A novel CSP ortholog group, CSP20, was also identified, likely missed in previous annotations because of low expression levels. Finally, CSP4 displayed evidence of repeated duplications but did not form a well-supported monophyletic group, preventing the unambiguous definition of paralogs, and included the sequence “RproCSPnew” defined in Marchant et al. (2021). This pattern may reflect low phylogenetic resolution and/or a complex evolutionary history involving ancient duplications followed by lineage-specific expansions and losses within the *Rhodnius* genus.

**Figure 4:**
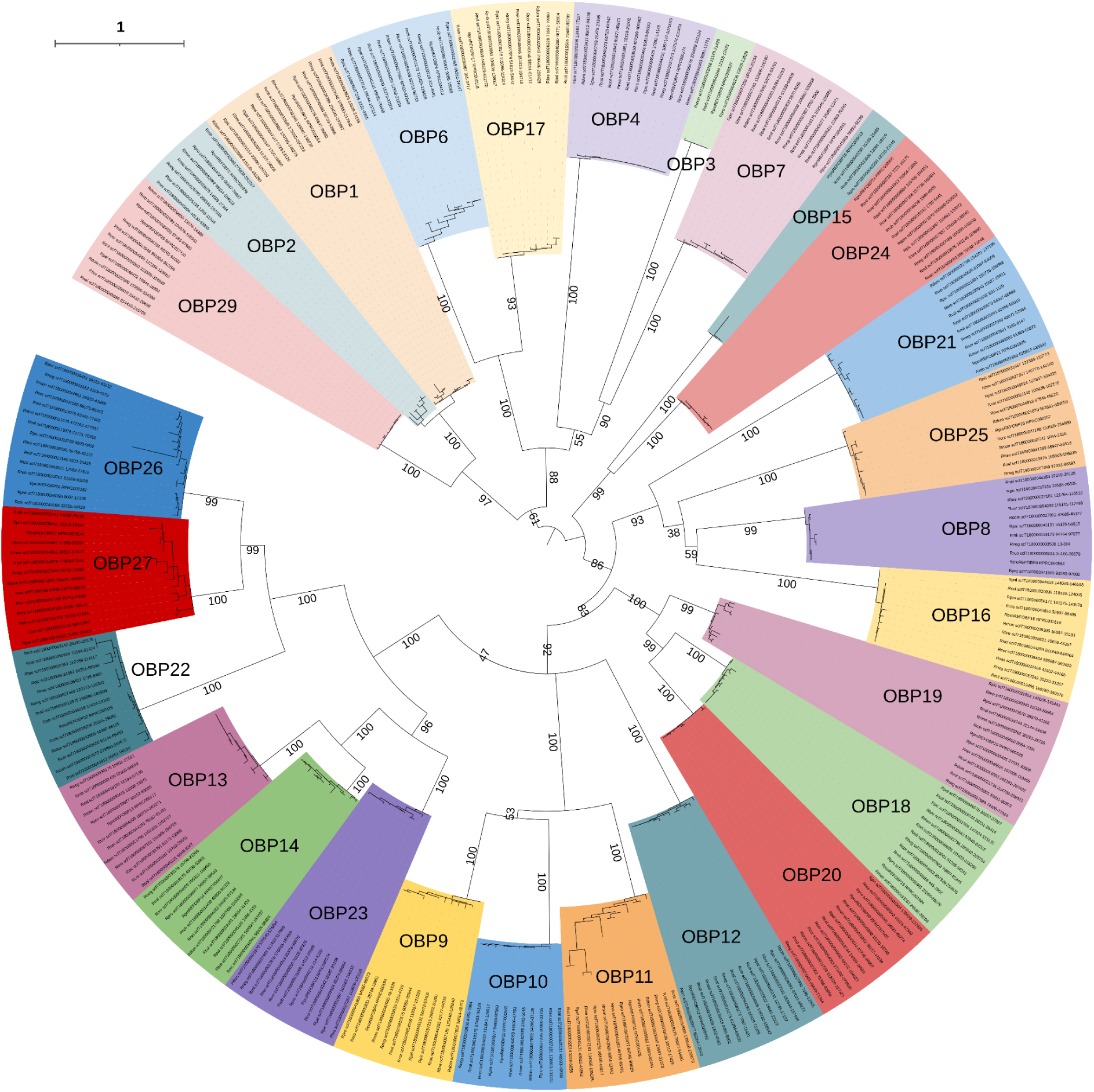
Consensus tree built from amino acid sequences of OBPs from 13 *Rhodnius* species. The phylogeny was built with IQ-TREE using the maximum likelihood approach. The model used is JTTDCMut+F+I+G4 chosen according to BIC. The bootstrap procedure was applied with 1,000 ultrafast bootstraps. OBPs are numbered according to Mesquita et al. (2015) and Volonté et al. (2025). Bootstrap values are indicated on branches except within defined orthologous groups, where they have been removed for clarity. The scale bar corresponds to 1 substitution per site.

**Figure 5:**
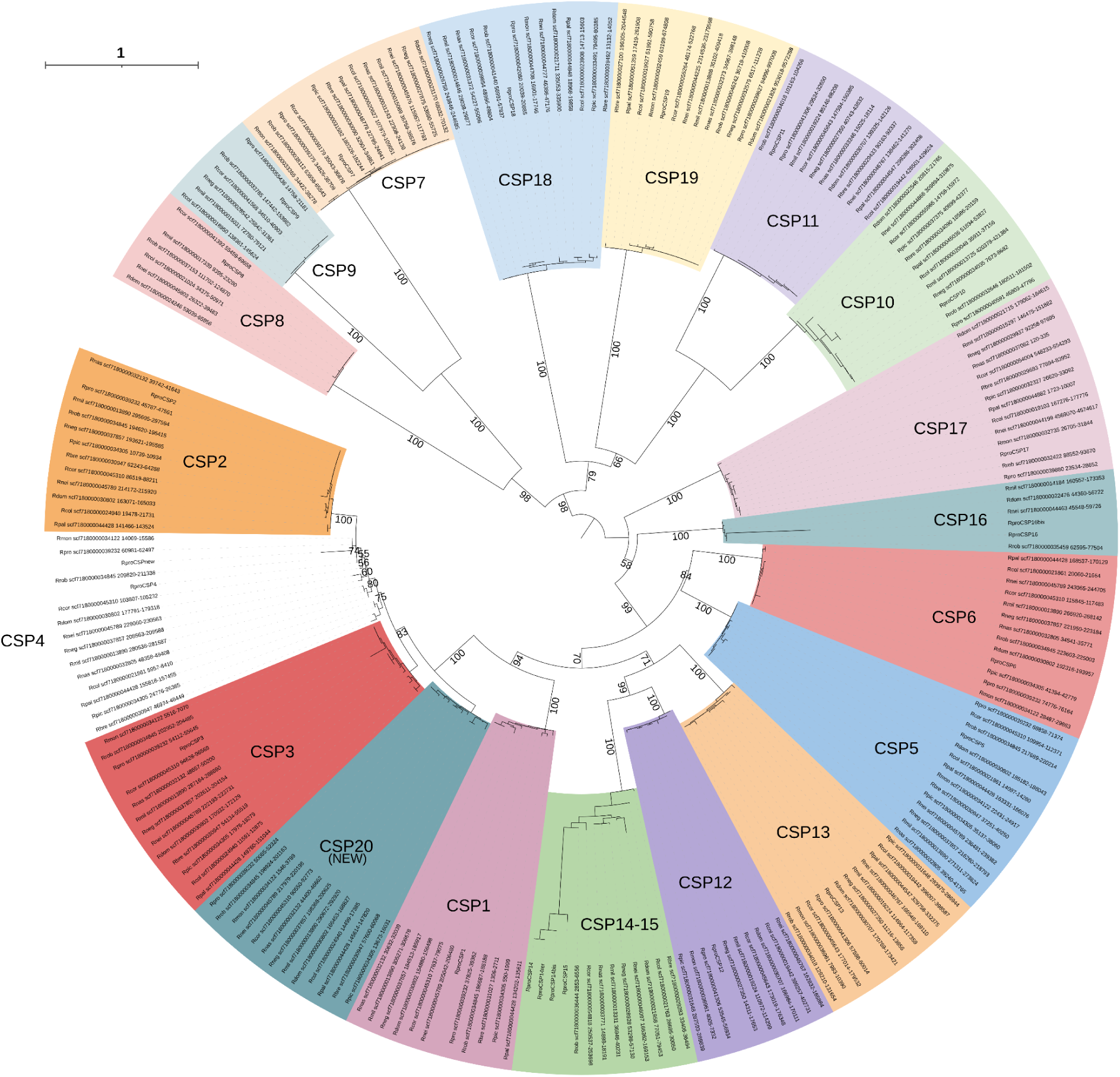
Consensus tree built from amino acid sequences of CSPs from 13 *Rhodnius* species. The phylogeny was built with IQ-TREE using the maximum likelihood approach. The model used is LG+G4 chosen according to BIC. The bootstrap procedure was applied with 1,000 ultrafast bootstraps. CSPs are numbered according to Latorre-Estivalis et al. (2017). Bootstrap values are indicated on branches except within defined orthologous groups, where they have been removed for clarity. We included the following sequences RproCSPnew, RproCSP14bis, RproCSP14ter and RproCSP16bis from Marchant et al. (2021). The scale bar corresponds to 1 substitution per site.

### Selection Pressure and Expression

Selection analyses were conducted on ortholog groups identified from phylogenetic reconstructions in order to investigate both ecological lineage-specific and site-specific evolutionary pressures acting on *Rhodnius* chemosensory genes. Analyses focused on the transmembrane chemosensory receptor gene families (ORs, GRs, and IRs), whereas the OBP and CSP gene families were excluded because their short sequences, typically encoding small soluble proteins of only 100–150 amino acids, substantially reduce the robustness and statistical power of dN/dS-based likelihood ratio tests. Branch models were used to test for differences in non-synonymous/synonymous substitution ratios (ω) between domiciliary and sylvatic lineages. Complete results, including M0 vs. M3 comparisons assessing heterogeneity in selective constraints among codon sites are provided in Supplementary Tables 8-10 and summary results are presented in Table 3. Because GR23 includes two previously described alternative transcripts that cannot be reliably distinguished by InsectOR, this gene was excluded from the selection analyses. In contrast, GR14 was retained because all sequences detected in the assemblies were unambiguously assigned to the GR14-RB transcript.

**Table 3:**
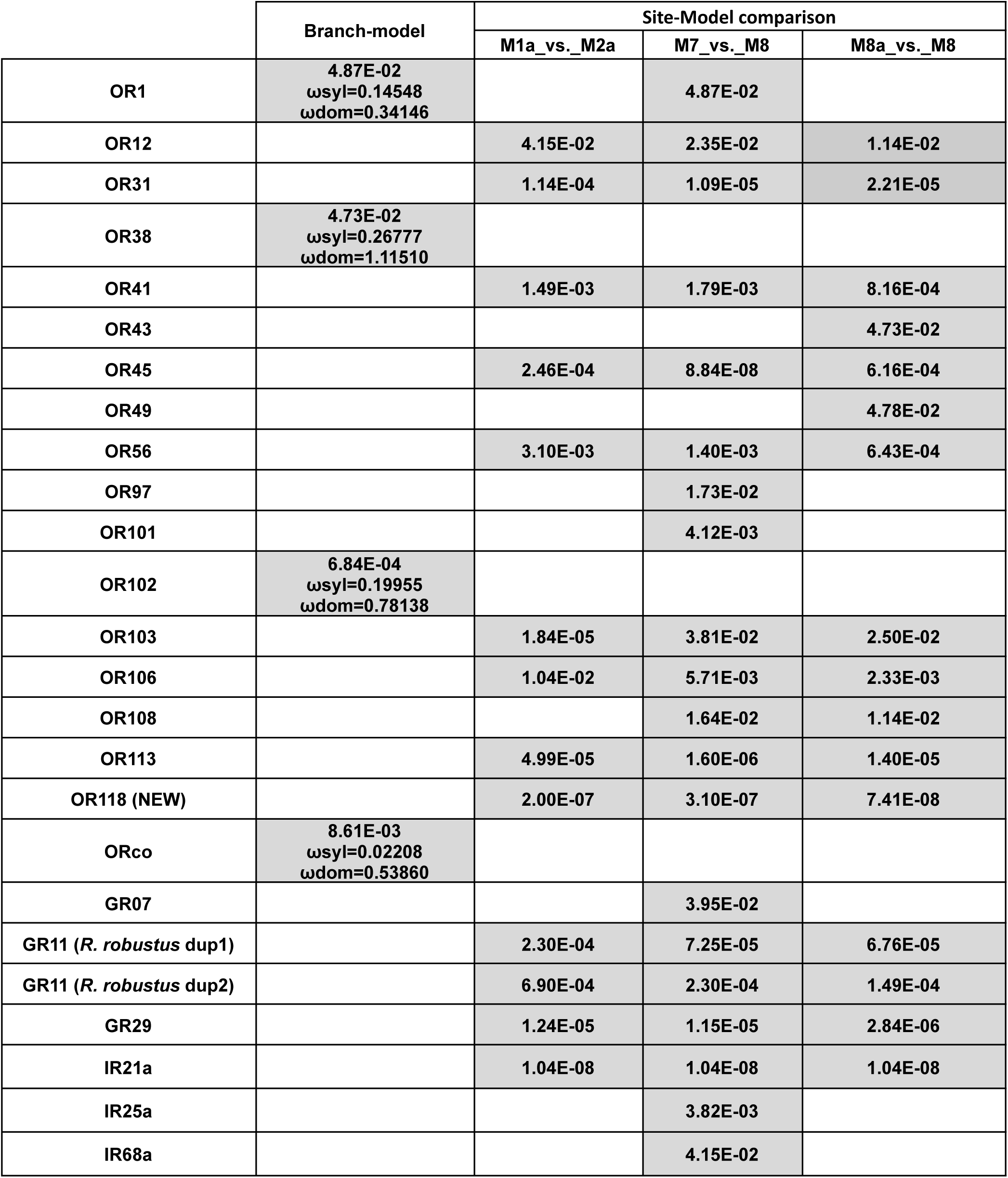
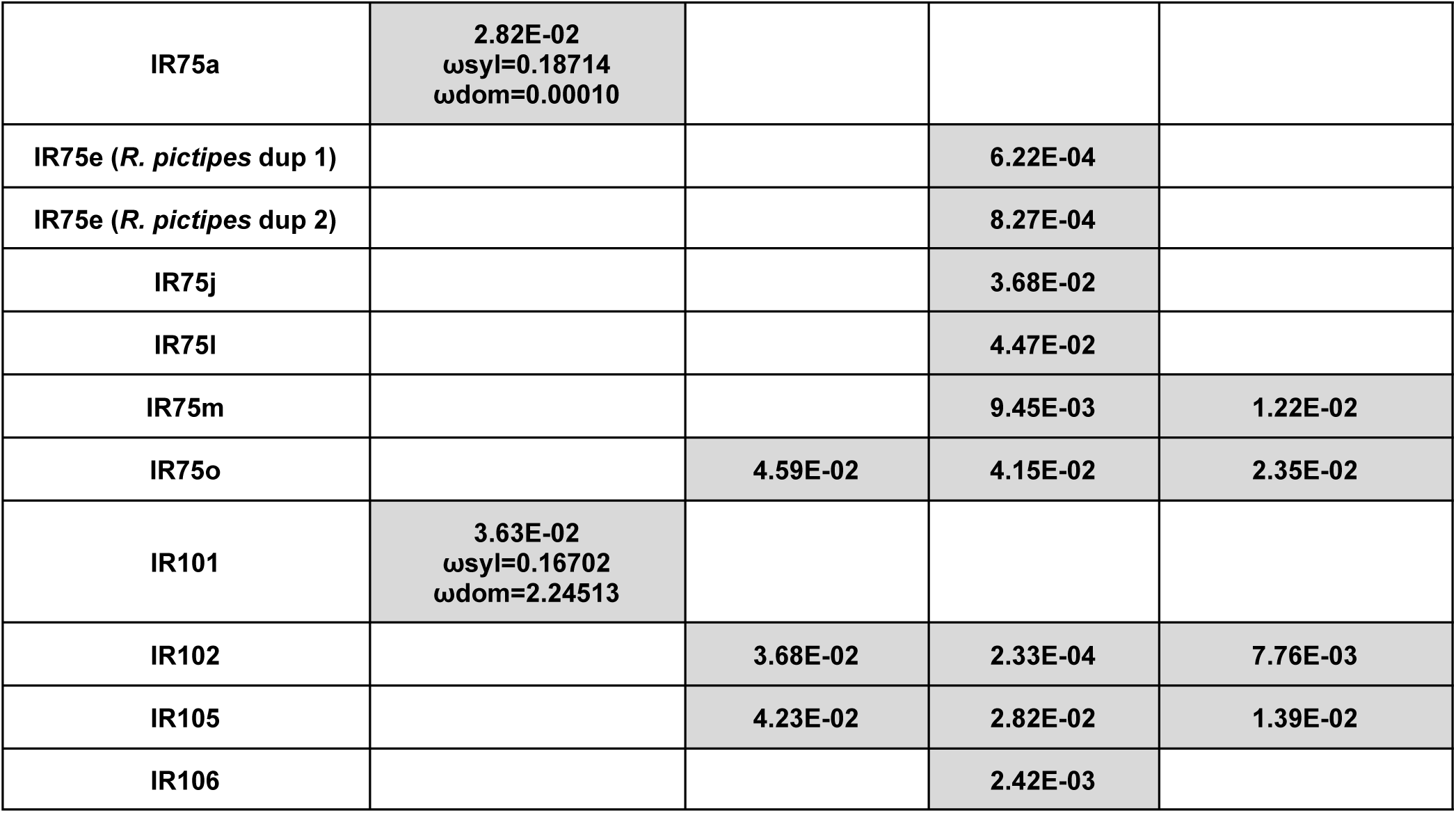
Chemoreceptors coding genes showing at least one significant result in CodeML comparisons. For each orthogroup and each test, the adjusted p-value (according to Benjamini and Hochberg) is indicated when lower than 5%.

|  | Branch-model | Site-Model comparison |  |  |
| --- | --- | --- | --- | --- |
|  |  | M1a_vs._M2a | M7_vs._M8 | M8a_vs._M8 |
| OR1 | 4.87E-02<br>$\omega_{\text{syl}}=0.14548$<br>$\omega_{\text{dom}}=0.34146$ | | 4.87E-02 | |
| OR12 |  | 4.15E-02 | 2.35E-02 | 1.14E-02 |
| OR31 |  | 1.14E-04 | 1.09E-05 | 2.21E-05 |
| OR38 | 4.73E-02<br>$\omega_{\text{syl}}=0.26777$<br>$\omega_{\text{dom}}=1.11510$ | | | |
| OR41 |  | 1.49E-03 | 1.79E-03 | 8.16E-04 |
| OR43 |  |  |  | 4.73E-02 |
| OR45 |  | 2.46E-04 | 8.84E-08 | 6.16E-04 |
| OR49 |  |  |  | 4.78E-02 |
| OR56 |  | 3.10E-03 | 1.40E-03 | 6.43E-04 |
| OR97 |  |  | 1.73E-02 |  |
| OR101 |  |  | 4.12E-03 |  |
| OR102 | 6.84E-04<br>$\omega_{\text{syl}}=0.19955$<br>$\omega_{\text{dom}}=0.78138$ | | | |
| OR103 |  | 1.84E-05 | 3.81E-02 | 2.50E-02 |
| OR106 |  | 1.04E-02 | 5.71E-03 | 2.33E-03 |
| OR108 |  |  | 1.64E-02 | 1.14E-02 |
| OR113 |  | 4.99E-05 | 1.60E-06 | 1.40E-05 |
| OR118 (NEW) |  | 2.00E-07 | 3.10E-07 | 7.41E-08 |
| ORco | 8.61E-03<br>$\omega_{\text{syl}}=0.02208$<br>$\omega_{\text{dom}}=0.53860$ | | | |
| GR07 |  |  | 3.95E-02 |  |
| GR11 (*R. robustus* dup1) |  | 2.30E-04 | 7.25E-05 | 6.76E-05 |
| GR11 (*R. robustus* dup2) |  | 6.90E-04 | 2.30E-04 | 1.49E-04 |
| GR29 |  | 1.24E-05 | 1.15E-05 | 2.84E-06 |
| IR21a |  | 1.04E-08 | 1.04E-08 | 1.04E-08 |
| IR25a |  |  | 3.82E-03 |  |
| IR68a |  |  | 4.15E-02 |  |
| IR75a | 2.82E-02<br>$\omega_{syl}=0.18714$<br>$\omega_{dom}=0.00010$ | | | |
| IR75e (*R. pictipes* dup 1) |  |  | 6.22E-04 |  |
| IR75e (*R. pictipes* dup 2) |  |  | 8.27E-04 |  |
| IR75j |  |  | 3.68E-02 |  |
| IR75l |  |  | 4.47E-02 |  |
| IR75m |  |  | 9.45E-03 | 1.22E-02 |
| IR75o |  | 4.59E-02 | 4.15E-02 | 2.35E-02 |
| IR101 | 3.63E-02<br>$\omega_{syl}=0.16702$<br>$\omega_{dom}=2.24513$ | | | |
| IR102 |  | 3.68E-02 | 2.33E-04 | 7.76E-03 |
| IR105 |  | 4.23E-02 | 2.82E-02 | 1.39E-02 |
| IR106 |  |  | 2.42E-03 |  |

Overall, branch-model analyses revealed contrasted evolutionary dynamics between sylvatic and domiciliary lineages. Four ORs and one IR exhibited higher ω ratios in domiciliary species, including Orco, OR1, OR38, OR102, and IR101. These increased ω ratios suggest a relaxation of purifying selection in domiciliary lineages and, for OR38 and IR101 where ω exceeded 1, possible adaptive divergence or accelerated evolution associated with domestic environments. The markedly higher ω ratio observed for Orco in domiciliary species also suggests substantial relaxation of selective constraints affecting this highly conserved co-receptor. In contrast, IR75a exhibited a markedly lower ω ratio in domiciliary lineages, indicating particularly strong purifying selection in species adapted to human dwellings. Together, these results suggest that the transition toward domiciliary habitats modified selective constraints acting on several chemosensory receptors. Among the genes showing no differential selective pressure between sylvatic and domiciliary species, some exhibit very low dN/dS ratios, indicating strong purifying selection (below 0.1). This includes 9 genes: OR2, GR1, GR2, GR15, IR25a, IR40a, IR68a, IR75h, and IR93a (Supplementary Tables 8-10).

Site-model analyses, which test for variation in ω among codon sites along sequences, revealed widespread signatures of positive selection in OR, GR, and IR genes. The M0 vs. M3 tests yielded significant results for almost all tested genes, except for 11 out of 101 tested (Supplementary Table 8-10). This indicates that these genes do not have a uniform evolutionary rate, but rather exhibit site-specific selective pressures. To further investigate, we performed additional tests to detect positive selection or ecological adaptation in the species.

Significant signals across all three site-model comparisons (M1a vs. M2a, M7 vs. M8, and M8a vs. M8) were detected for 9 ORs (OR12, OR31, OR41, OR45, OR56, OR103, OR106, OR113, and the newly identified OR118), for 3 GRs (both duplicates of GR11 and GR29), and for 5 IRs (IR102, IR105, IR75m, IR75o and IR21a) indicating robust evidence of positive selection acting on these genes. Among them, IR21a showed the strongest signal (padj = 1.04 × 10⁻⁸). Additional weaker but significant signals, supported by only one or two comparisons, were detected for 6 ORs (OR1, OR43, OR49, OR97, OR101, and OR108), one GR (GR7) and 8 IRs (IR25a, IR68a, the two duplicates of IR75e, IR75j, IR75l, IR75m and IR106). The combination of branch-model and site-model signals for OR1 further supports ongoing adaptive divergence in domiciliary lineages.

Relative transcript abundance among the 9 species with gene expression data of the 6 genes displaying significant differences in ω ratios between ecological groups is shown in Figure 6. The three species with the highest transcription levels for Orco are, in descending order: *R. pallescens*, *R. nasutus*, and *R. neglectus*, all of which are among the 4 domiciliary species. Similarly, for the gene OR38, the top three species are *R. nasutus*, *R. neglectus*, and *R. prolixus*, again all domiciliary species.

**Figure 6:**
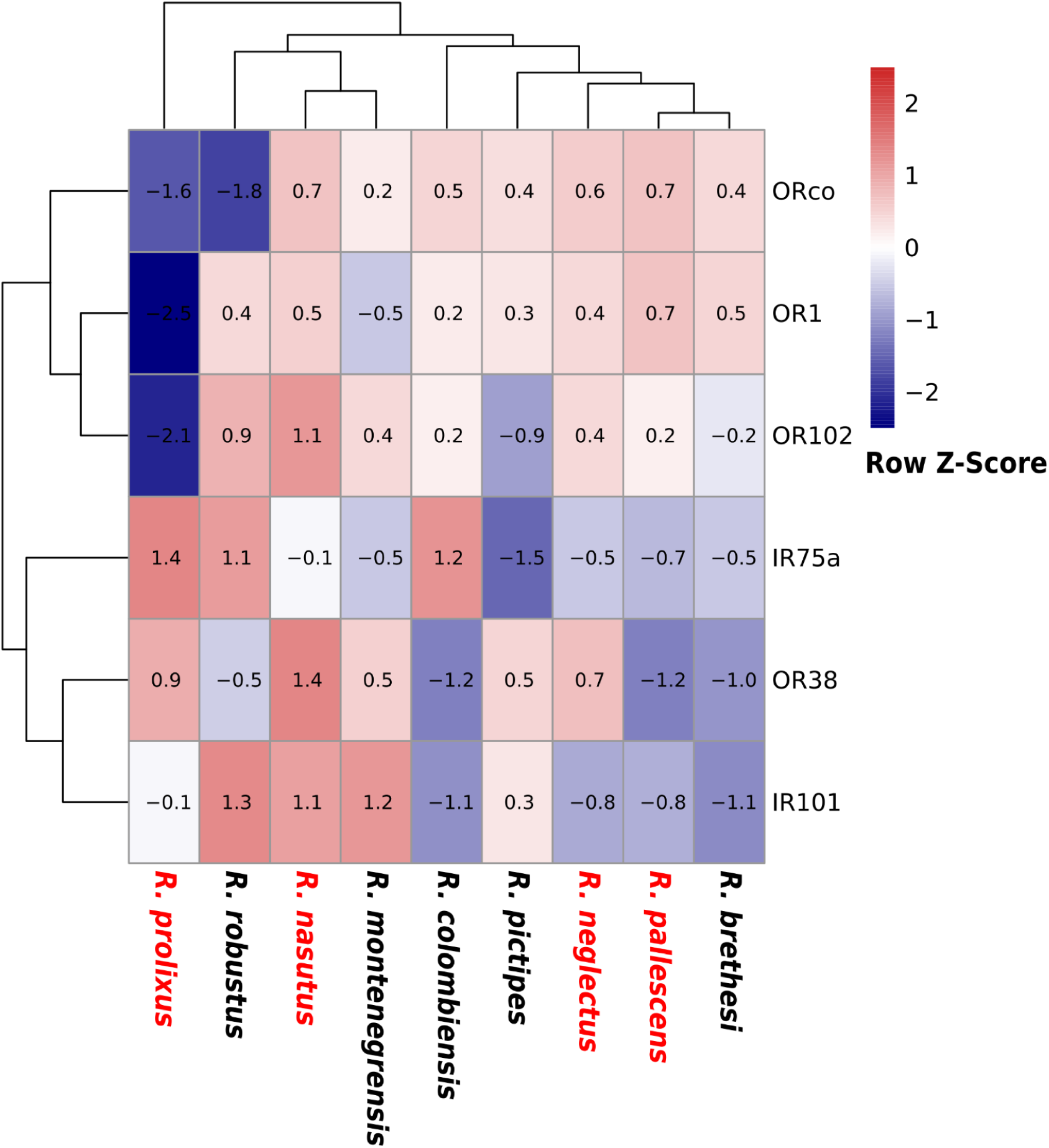
Heatmap of relative transcript abundance for genes showing differential dN/dS ratio between domiciliary species and sylvatic species. Transcript expression levels were normalized as log2(TPM+1) and scaled by row (row Z-score). The color scale represents relative expression levels: red indicates over-expression relative to the gene’s mean expression, while blue indicates under-expression. Hierarchical clustering was performed on rows and columns. Domiciliary species are indicated in red.

## Discussion

Chemosensory systems are expected to evolve under strong ecological constraints because they mediate host detection, feeding behavior, and habitat selection. Using comparative analyses across 13 *Rhodnius* species, our results provide insight into how different chemosensory gene families evolved within a hematophagous lineage characterized by ecological diversification and repeated transitions toward domiciliary habitats.

### Conservation and Diversification of Chemosensory Repertoires in Rhodnius

A large proportion of chemosensory receptors remained conserved across *Rhodnius* species, especially for GRs and IRs, for which most ortholog groups followed a 1:1 orthology pattern and displayed limited sequence divergence. These observations suggest that a substantial part of the chemosensory repertoire was already established in the common ancestor of this hematophagous genus and remained globally conserved during diversification of extant *Rhodnius* species. Twenty-eight GRs were annotated in *Rhodnius* species, a number comparable to that observed in the hematophagous bed bug *Cimex lectularius* (35 GRs; Hansen et al. 2014), but markedly lower than those found in non-hematophagous hemipterans such as the phytophagous aphid *Acyrthosiphon pisum* (77 GRs; Smadja et al. 2009) or *Oncopeltus fasciatus* (over 150 GRs; Panfilio et al. 2019). This reduced GR repertoire is consistent with a specialized blood-feeding diet and may be associated with a narrower range of gustatory stimuli linked to hematophagy.

In contrast, OR repertoires displayed stronger evolutionary dynamics. Considering Hemiptera, the *Rhodnius* OR repertoire (123 ORs, including Orco) is substantially larger than that of the non-triatomine hematophagous bed bug *Cimex lectularius* (50 ORs; Hansen et al. 2014) and more similar in size to those reported in the seed-feeding *Oncopeltus fasciatus* (122 ORs; Panfilio et al. 2019), the polyphagous *Halyomorpha halys* (138 ORs; Sun et al. 2020), and the predatory *Gerris buenoi* (155 ORs; Armisén et al. 2018). This suggests that the transition to blood-feeding in Triatominae did not result in the marked OR repertoire reduction as observed in the highly specialized hematophagous bed bug *Cimex lectularius*.

The strong diversification observed in *Rhodnius* ORs contrasted with the overall conservation of GR, IR, OBP and CSP repertoires. Similar differences among receptor families have been reported in other insects, where OR repertoires often evolve more rapidly than GRs and conserved IRs in relation to ecological divergence (McBride 2007; Smadja et al. 2012). Transcriptomic analyses previously conducted in *Rhodnius* also revealed relatively stable expression profiles for GR and IR proteins across developmental stages (Latorre-Estivalis et al. 2022), supporting comparatively constrained evolution for these gene families relative to OR repertoires. Specifically, Latorre-Estivalis et al. (2022) identified several age-dependent expression profiles among ORs, including four early-decreasing ORs (OR17, OR39, OR96, OR104), a late-decreasing OR (OR100), and 13 ORs displaying increased expression with age. Together, these observations support greater regulatory flexibility in ORs compared with other chemosensory receptor families.

Several OR clades, including OR21-23, OR28-29, and OR54, showed lineage-specific expansions associated with tandem duplications potentially linked to functional diversification. The OR28-29 group was particularly expanded, with 49 genes identified across the 13 assemblies. In addition, ORs represented the receptor family with the largest number of genes showing signatures of positive selection at the site level, with 11 ORs displaying significant results for at least two models, including M8a vs. M8, the most stringent test. However, signatures of divergent evolution were also detected in 3 GRs and 5 IRs. These results suggest that in *Rhodnius*, the ORs constitute the most evolutionarily dynamic component of the chemosensory repertoire, the acquisition of new paralogs potentially providing novel sensitivities required for the complex chemical landscape of vertebrate hosts. Moreover, the evidence of recurrent site-specific positive selection across the evolutionary history of the ortholog group for both ORs, GRs and IRs suggests recurrent functional diversification of chemosensory receptors potentially associated with ecological diversification.

### Ecological Specialization

#### OR Diversification

The genus *Rhodnius* occupies ecological niches ranging from highly specialized palm-associated habitats to broader sylvatic ones and to domiciliary environments. Although genome completeness prevents robust quantitative comparisons of repertoire sizes between sylvatic and domiciliary species, repeated tandem duplications within OR21-23 and OR28-29 are particularly observed in the two sylvatic species *R. colombiensis* and *R. robustus*.

These marked OR expansions may reflect distinct but convergent ecological contexts associated with recent diversification processes within *Rhodnius*. For *R. robustus,* a wide Amazonian distribution and broad range of sylvatic habitats associated with multiple palm species and bromeliads are described (Abad-Franch et al. 2005). Molecular studies revealed several cryptic lineages in *R. robustus,* suggesting a relatively recent Amazonian radiation with limited divergence among members of the *robustus* complex (Filée et al. 2022). Similarly, *R. colombiensis,* endemic to central Colombia, mainly reported from the upper Magdalena River basin, belongs to the recently diversified trans-Andean *pallescens* group and occupies heterogeneous ecological environments associated with *Attalea* palm trees at the sylvatic–peridomestic interface (Díaz et al. 2014; Filée et al. 2022; Hernández et al. 2025). It is interesting to note that *R. coreodes*, sometimes classified within the separate genus *Psammolestes* (which includes three species, with similar ecologies), does not exhibit a restricted chemosensory gene repertoire, despite having a narrow host range limited to birds, mainly from the Furnariidae and Psittacidae families (Hernández et al. 2020).

Overall, the combination of lineage-specific OR expansions, signatures of positive selection, and dynamic transcriptomic regulation supports greater interspecific divergence within the OR family compared with other chemosensory groups.

#### Diversification of conserved ionotropic pathways

Unexpected diversification emerged within the IR family despite the overall conservation of ionotropic repertoires. Repeated duplications of the conserved co-receptor IR8a were identified in several species, especially *R. colombiensis*. Because IR8a is generally maintained as a single-copy gene in insects, these duplications suggest an unusual evolutionary trajectory within *Rhodnius*. Although assembly artifacts cannot be fully excluded, the recurrence of these duplications across assemblies supports their biological relevance. Additional diversification was detected within the IR75 clade, including positive selection affecting IR75m and IR75o and duplicated IR75e copies in *R. pictipes*. These observations indicate that some conserved ionotropic pathways also underwent lineage-specific diversification, although to a lower extent than OR repertoires.

#### Functional Diversification of Bitter-Associated GRs

Although GR repertoires remained globally conserved, some receptors displayed evidence of functional diversification.

Among other gustatory receptors, GR5 and GR11 have been proposed to be involved in bitter compound detection based on homology with functionally characterized receptors (Mesquita et al. 2015). Bitter compounds are commonly associated with toxic or harmful substances and often act as warning signals in both plants and animals. The signatures of positive selection detected in GR11 therefore suggest recurrent functional diversification of bitter perception during *Rhodnius* evolution, potentially in response to diverse environmental toxic compounds encountered across sylvatic and domestic habitats. Although site-model analyses do not support lineage-specific interpretations, exposure to insecticides in domiciliary environments may also contribute to shaping bitter compound perception in some species.

#### Conservation of Sugar Detection Pathways

GR1 clustered with fructose receptors previously characterized in *Drosophila melanogaster* and *Apis mellifera* (Sato, Tanaka, and Touhara 2011; Takada et al. 2018) and remained highly conserved across species, as evidenced by its low dN/dS ratio (dN/dS = 0.09125; see Supplementary Table 9). This GR1 is furthermore highly expressed in the antenna of *R. prolixus* adults (Latorre-Estivalis et al. 2017) and in our study, shows a high expression in the nine *Rhodnius* species included in the expression heatmap. Such conserved and elevated expression levels are somewhat surprising in hematophagous insects and suggest the maintenance of sugar detection capacities in Triatominae. Interestingly, it has been shown that *R. prolixus* could be fed on sucrose solution or even on tomato, behaviors that subsequently increased blood meal intake and improved fitness-related traits, including survival and lifespan (Díaz-Albiter et al. 2016). In sylvatic habitats, *Rhodnius* species are closely associated with palm trees, where sugar resources may be available from sap or fruits. Consistent with this hypothesis, palm DNA was recently detected in the gut contents of *R. robustus*, suggesting occasional plant feeding in natural conditions (Da Lage et al. 2024). Together, these observations support the hypothesis that sugar perception remains functionally relevant in Triatominae despite their hematophagy. *Rhodnius* species may therefore have retained part of the sugar metabolic pathway and sugar detection capacities inherited from their phytophagous ancestors, from which hematophagous Hemiptera diverged approximately 230 million years ago (Wootton 1981).

#### Absence of Canonical CO₂-associated GRs

Despite the major role of carbon dioxide in triatomine host-seeking behaviour and its importance as a long-range cue for vertebrate host detection (Lazzari, Pereira, and Lorenzo 2013; Guerenstein and Lazzari 2009), we found no evidence of the gustatory receptors classically involved in CO₂ detection in other insects, corroborating the findings of Mesquita et al. (2015). This suggests that CO₂ sensing in Triatominae may rely on alternative molecular mechanisms. In particular, CO₂ detection could be mediated by other receptor families such as IRs. Interestingly, studies in *Drosophila* showed that attraction to CO₂ during foraging can involve an alternative IR25a-dependent pathway independent of the GR pathway (van Breugel, Huda, and Dickinson 2018; Yan et al. 2020). These observations suggest that conserved behavioral functions may rely on distinct receptor systems in different insect lineages.

#### Functional Versatility of OBPs and CSPs

Although OBP and CSP repertoires remained globally conserved across *Rhodnius* species, increasing evidence suggests that these soluble proteins may fulfill functions extending far beyond odorant transport. Transcriptomic analyses revealed strong developmental upregulation of many CSPs and OBPs during adult maturation (Latorre-Estivalis et al. 2022), consistent with their involvement in adult-specific sensory functions such as host seeking, mating, and oviposition. Functional studies in *R. prolixus* further support specialized roles for some OBPs in pheromone perception and reproductive behaviours (Oliveira et al. 2018). Beyond chemosensation, OBPs and CSPs have increasingly been implicated in detoxification and responses to chemical stress in insects (Wang et al. 2025). Although such functions remain unexplored in *Rhodnius*, the strong conservation of several OBP and CSP ortholog groups across species suggests that these proteins may contribute to physiological processes essential for adaptation to diverse environmental conditions. This functional versatility makes OBPs and CSPs promising targets for future studies aiming to interfere with both chemosensory behaviors and adaptive physiological responses in triatomine vectors.

Together, these results suggest that ORs constitute the most dynamic component of the *Rhodnius* chemosensory repertoire and may represent major contributors to sensory diversification and ecological adaptation within the genus.

### Domiciliation and Remodeling of Selective Pressures

If the overall repertoire sizes remained globally conserved between sylvatic and domiciliary species, several receptors showed differential selective pressures between habitats (Orco, OR1, OR38, OR103, IR75a, and IR101), including two (OR38 and IR101) showing a high positive selection in domiciliary species and one (OR1) exhibiting signatures of site-specific positive selection. Orco and OR38 showed a very high expression in domiciliary species.

Because domiciliation occurred independently across multiple *Rhodnius* lineages, these repeated shifts in selective pressures, two of which are supported by higher transcriptomic expression in domiciliary species, are consistent with convergent adaptation associated with domiciliary environments.

The signal detected for Orco was particularly notable. Orco is a highly conserved olfactory co-receptor required for OR function across insects and is generally maintained under strong purifying selection. Functional studies in *Aedes aegypti*, *Anopheles gambiae*, *Manduca sexta*, and *Cochliomyia hominivorax* showed that disruption of Orco strongly impairs host-seeking and feeding behaviors (DeGennaro et al. 2013; Fandino et al. 2019; Paulo et al. 2021). In *R. prolixus*, RNAi-mediated silencing of Orco disrupts host finding, blood feeding, oviposition, molting, and survival (Franco et al. 2016). Against this conserved functional background, the elevated ω values observed in domiciliary lineages suggest shifts in selective constraints acting on a central component of the olfactory system. Unlike lineage-specific OR expansions, the evolutionary signal detected in Orco may reflect changes affecting global olfactory regulation rather than diversification of individual odorant receptors.

## Conclusion and Implications for Vector Control

The contrasting evolutionary dynamics observed among chemosensory families have implications for vector adaptation. OR repertoires displayed the strongest lineage-specific expansions and signatures of positive selection, whereas GRs, IRs, OBPs, and CSPs remained comparatively conserved across species despite evidence of functional diversification in some specific lineages or ortholog groups.

The strong interspecific divergence observed in OR repertoires suggests that interventions targeting OR-mediated pathways may require species-specific optimization. Conversely, the higher conservation of GRs, IRs, OBPs, and CSPs may provide more stable targets for broad-spectrum approaches potentially effective across multiple triatomine vectors. Especially, the functional versatility of OBPs and CSPs makes them promising targets for future studies aiming to interfere with both chemosensory behaviours and adaptive physiological responses in triatomine vectors.

## Abbreviations

CSP: chemosensory protein
GR: gustatory receptor
IR: ionotropic receptor
OBP: odorant-binding protein
OR: odorant receptor
TPM: transcripts per million

## Data Availability

All identified chemosensory genes in both amino acid and nucleotide, multiple sequence alignments, the resulting phylogenetic trees and the results of selection pressure analyses are available at 10.5281/zenodo.19064793. All other datasets related to genome assemblies are available through the associated Zenodo community [KissingBugsOmics](https://zenodo.org/communities/kissingbugsomics/) and the NCBI BioProject [PRJNA1405103](https://www.ncbi.nlm.nih.gov/bioproject/1405103).

## Acknowledgments

We would like to thank all our collaborators who provided us with the new CTA strains used in this study: João Aristeu da Rosa and Jader Oliveira (Universidade Estadual Paulista (Unesp), Faculdade de Ciências Farmacêuticas, Araraquara, São Paulo, Brazil). Thanks to Claire Capdevielle-Dulac for performing the DNA extractions that led to the genomic assemblies. We acknowledge the GenOuest bioinformatics core facility (https://www.genouest.org) for providing the computing infrastructure.

---

**Supplementary Figure 1:**
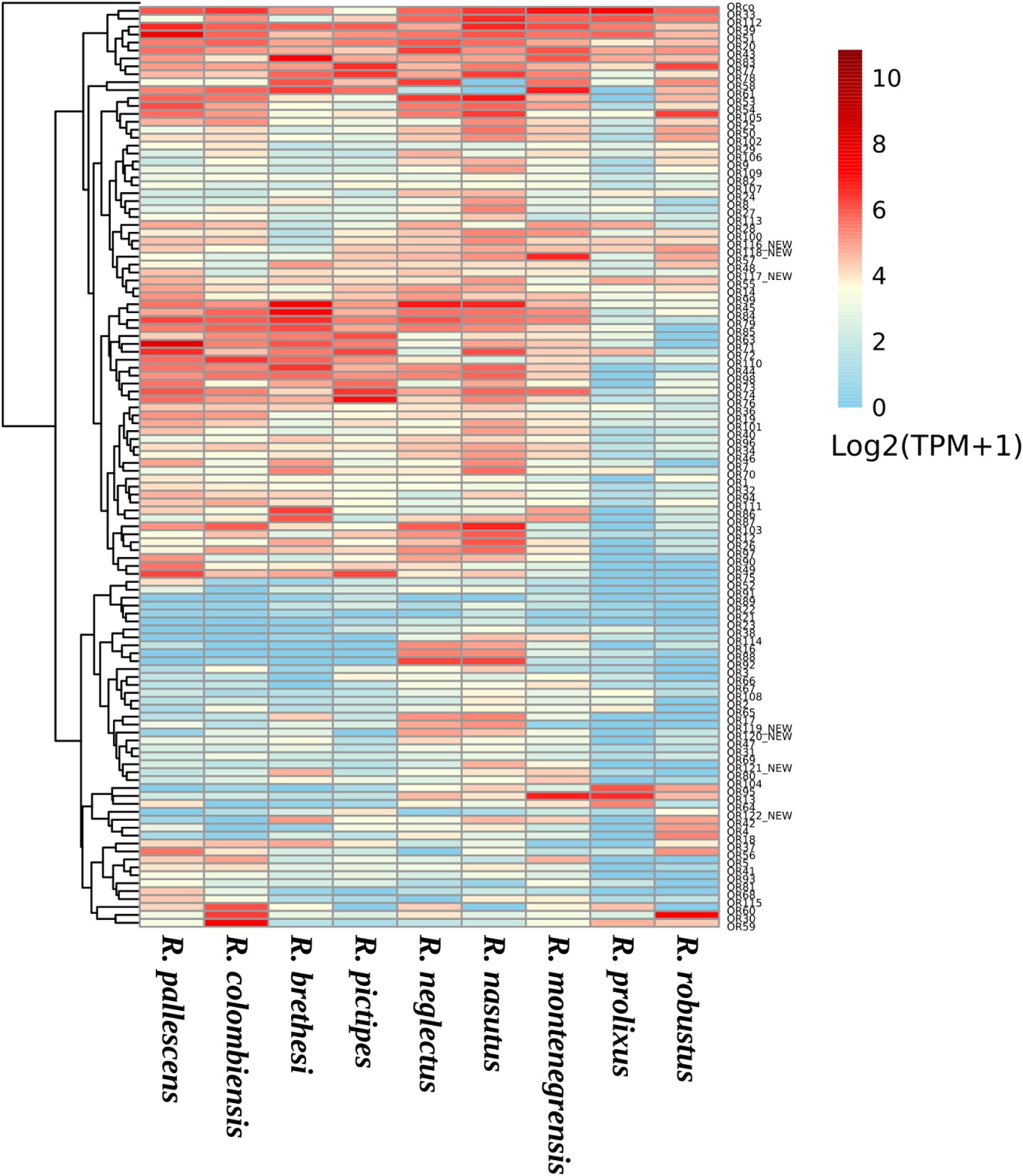
TPM-based Expression Profiles of *Rhodnius* ORs Genes. Data are expressed as log2(TPM+1) values. TPM (Transcripts Per Million) were calculated and normalized for both sequencing depth and gene length. Hierarchical clustering was applied to rows to group genes with similar expression patterns.

**Supplementary Figure 2:**
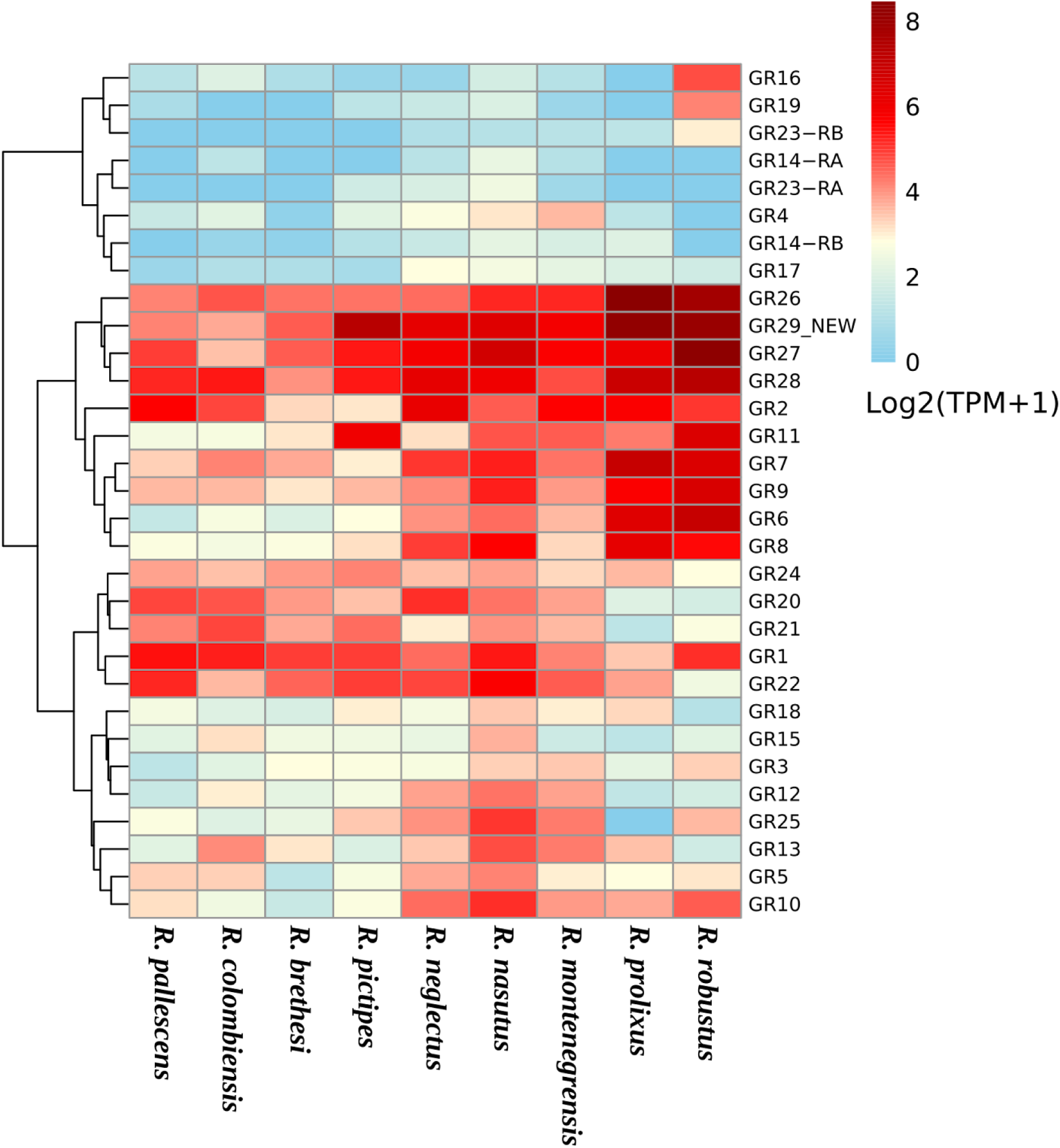
TPM-based Expression Profiles of *Rhodnius* GRs Genes. Data are expressed as log2(TPM+1) values. TPM (Transcripts Per Million) were calculated and normalized for both sequencing depth and gene length. Hierarchical clustering was applied to rows to group genes with similar expression patterns.

**Supplementary Figure 3:**
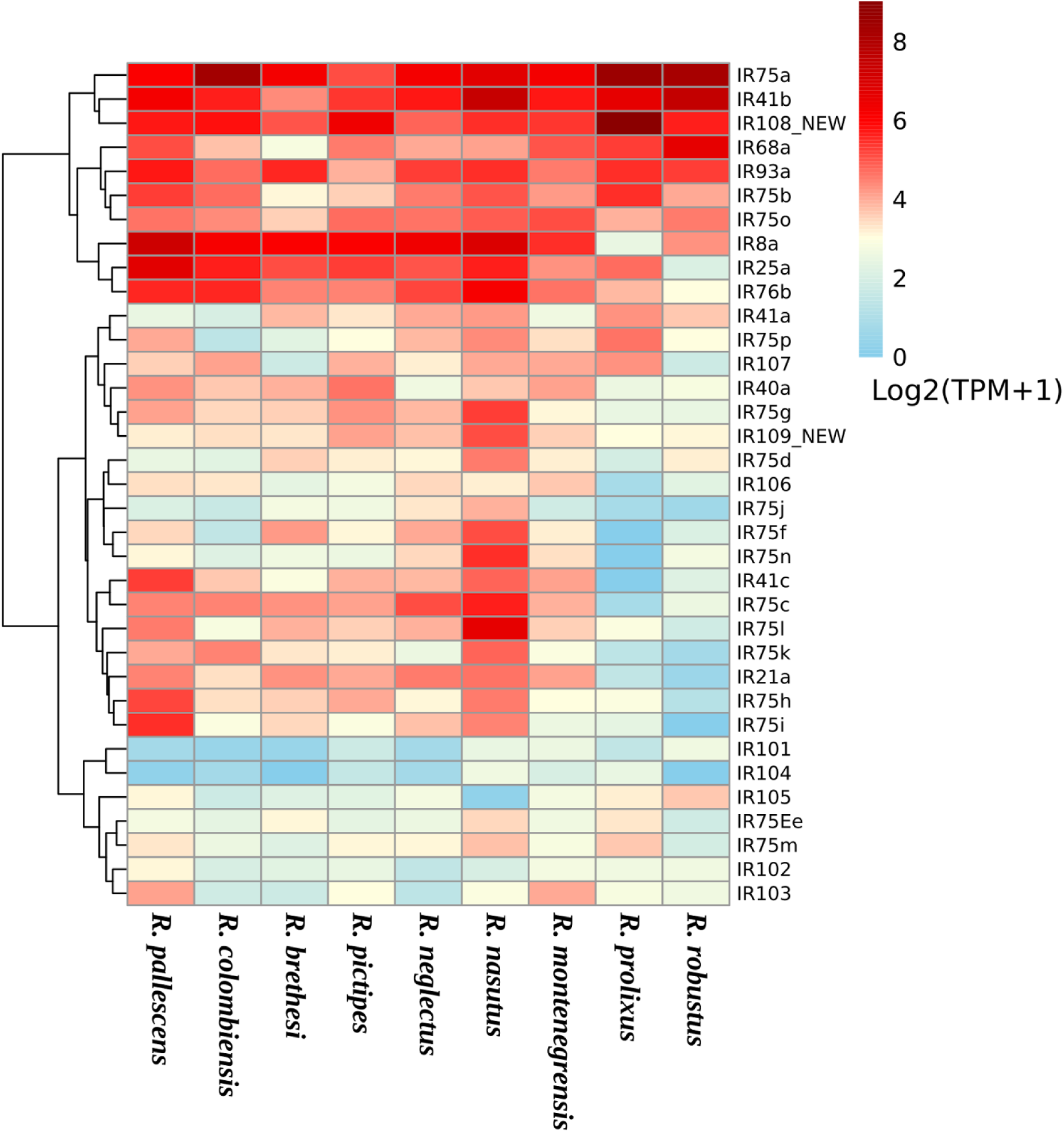
TPM-based Expression Profiles of *Rhodnius* IRs Genes. Data are expressed as log2(TPM+1) values. TPM (Transcripts Per Million) were calculated and normalized for both sequencing depth and gene length. Hierarchical clustering was applied to rows to group genes with similar expression patterns.

**Supplementary Figure 4:**
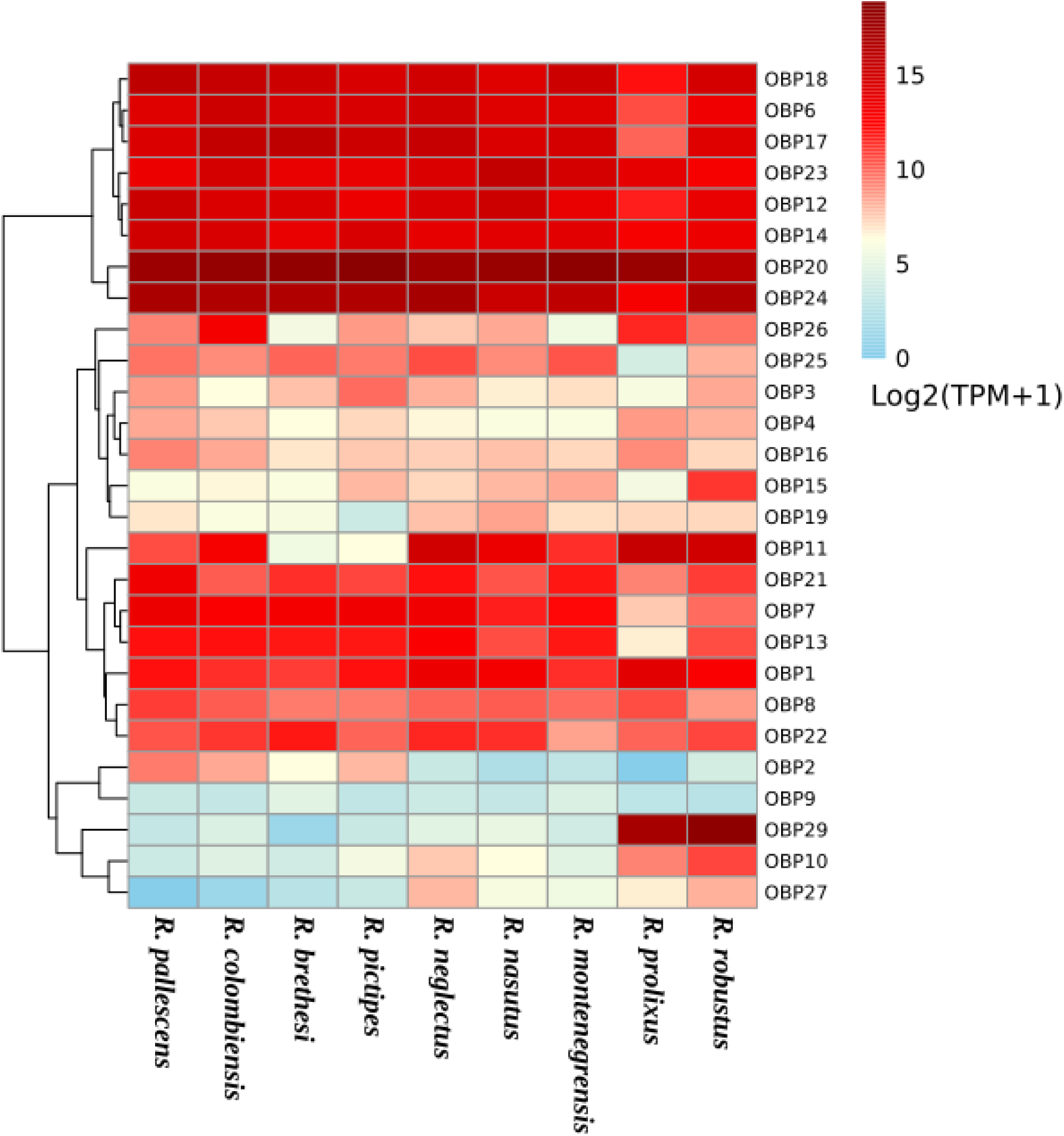
TPM-based Expression Profiles of *Rhodnius* OBPs Genes. Data are expressed as log2(TPM+1) values. TPM (Transcripts Per Million) were calculated and normalized for both sequencing depth and gene length. Hierarchical clustering was applied to rows to group genes with similar expression patterns.

**Supplementary Figure 5:**
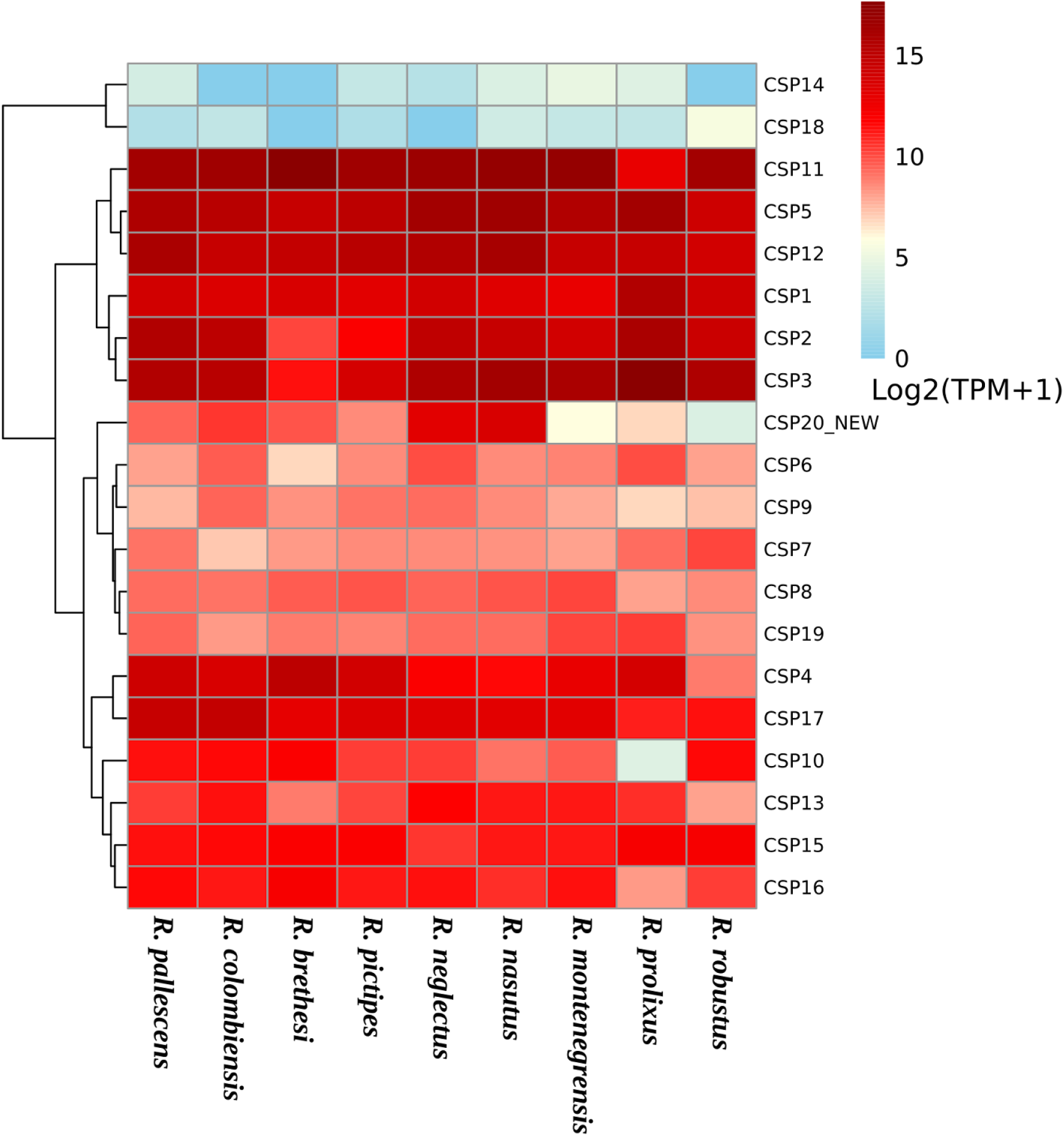
TPM-based Expression Profiles of *Rhodnius* CSPs Genes. Data are expressed as log2(TPM+1) values. TPM (Transcripts Per Million) were calculated and normalized for both sequencing depth and gene length. Hierarchical clustering was applied to rows to group genes with similar expression patterns.

**Supplementary Table 1:** Summary of Hemipteran query sequences used for chemosensory gene annotation. This table details the reference sequences (queries) utilized for the homology-based annotation of the chemosensory repertoire in this study. Empty reference fields indicate sequences derived directly from NCBI automatic annotation pipelines.

| Family | Species | OBPs | CSPs | ORs | GRs | IRs | References |
| --- | --- | --- | --- | --- | --- | --- | --- |
| Aphididae | *Acyrtosiphon pisum* | 13 | 10 | 79 | 77 | - | Smadja et al. 2009 |
| Aphididae | *Aphis gossypii* | 9 | 9 | - | - | - | - |
| Aphididae | *Myzus persicae* | 9 | 9 | - | - | - | - |
| Aphididae | *Sitobion avenae* | 11 | 7 | - | - | - | - |
| Cimicidae | *Cimex lectularius* | 10 | 8 | 50 | 35 | 41 | Hansen et al. 2014 / Benoit et al. 2016 |
| Delphacidae | *Laodelphax striatellus* | 9 | 12 | - | - | - | - |
| Delphacidae | *Nilaparvata lugens* | 10 | 11 | - | - | - | - |
| Delphacidae | *Sogatella furcifera* | 12 | 9 | - | - | - | - |
| Miridae | *Adelphocoris lineolatus* | 34 | 14 | 88 | - | - | Xiao et al. 2016 |
| Miridae | *Adelphocoris suturalis* | 16 | 8 | - | - | - | - |
| Miridae | *Apolygus lucorum* | 38 | 8 | 110 | - | 9 | An et al. 2016 |
| Miridae | *Lygus lineolaris* | 33 | - | - | - | - | - |
| Miridae | *Cyrtorhinus lividipennis* | - | - | 15 | - | - | Wang et al. 2018 |
| Pentatomidae | *Nezara viridula* | 42 | 22 | - | - | - | - |
| Pentatomidae | *Halyomorpha halys* | - | - | 138 | - | 77 | Sun et al. 2020 / Sparks et al. 2020 |
| Reduviidae | *Rhodnius prolixus* | 27 | 18 | 111 | 28/30 | 33 | Mesquita et al. 2015 |
| Reduviidae | *Triatoma brasiliensis* | 24 | 16 | - | - | - | - |
| Reduviidae | *Triatoma infestans* | - | - | - | - | 47 | Latorre Estivalis et al. 2022 |
| Reduviidae | *Triatoma rubrofasciata* | 19 | - | - | - | - | - |
| Pediculidae | *Pediculus humanus* | 7 | 8 | - | 5 | - | NCBI |
| Gerridae | *Gerris buenoi* | - | - | 155 | - | - | Armisen et al. 2018 |
| Lygaeidae | *Nysius ericae* | - | - | 83 | - | - | Zhang et al. 2016 |
| Lygaeidae | *Oncopeltus fasciatus* | - | - | 122 | 115/169 | 37 | Panfilio et al. 2019 |
| Berytidae | *Yemma signatus* | - | - | 66 | - | - | Song et al. 2020 |
| Psyllidae | *Diaphorina citri* | - | - | 46 | - | - | Wu et al. 2016 |
| Psyllidae | *Cacopsylla chinensis* | - | - | 7 | - | - | Xu et al. 2019 |
| Pseudococcidae | *Phenacoccus solenopsis* | - | - | 4 | - | - | Nie et al. 2017 |

**Supplementary Table 2.** Putative chemoreceptor genes and pseudogenes annotated using InsectOR. Domiciliary species are indicated in red. The first column shows the number of complete, non-pseudogenized sequences with a start codon (>300 aa) retained for analysis. Additional columns include: raw counts of complete genes (>300 aa), partial genes (<300 aa), pseudogenes without frameshifts, pseudogenes with frameshifts, and total sequences. For IRs, counts are preliminary due to the need for manual curation.

| Species | Complete + start | >300 aa | <300 aa | Non-pseudogenes | Pseudogenes | Total |
| --- | --- | --- | --- | --- | --- | --- |
| **ORs** |  |  |  |  |  |  |
| *R. brethesi* | 66 | 98 | 29 | 121 | 6 | 127 |
| *R. colombiensis* | 78 | 114 | 27 | 132 | 9 | 141 |
| *R. coreodes* | 70 | 110 | 27 | 115 | 22 | 137 |
| *R. domesticus* | 72 | 105 | 42 | 132 | 15 | 147 |
| *R. milesi* | 82 | 121 | 31 | 138 | 14 | 152 |
| *R. montenegrensis* | 70 | 102 | 48 | 135 | 15 | 150 |
| *R. nasutus* | 34 | 59 | 39 | 92 | 6 | 98 |
| *R. neglectus* | 74 | 101 | 41 | 136 | 6 | 142 |
| *R. neivai* | 84 | 135 | 27 | 150 | 12 | 162 |
| *R. pallescens* | 24 | 49 | 45 | 91 | 3 | 94 |
| *R. pictipes* | 70 | 110 | 44 | 143 | 11 | 154 |
| *R. prolixus* | 69 | 99 | 47 | 131 | 15 | 146 |
| *R. robustus* | 83 | 120 | 62 | 166 | 16 | 182 |
| **GRs** |  |  |  |  |  |  |
| *R. brethesi* | 25 | 29 | 10 | 35 | 4 | 39 |
| *R. colombiensis* | 28 | 28 | 6 | 32 | 2 | 34 |
| *R. coreodes* | 23 | 28 | 8 | 31 | 5 | 36 |
| *R. domesticus* | 28 | 31 | 3 | 32 | 2 | 34 |
| *R. milesi* | 28 | 29 | 6 | 34 | 1 | 35 |
| *R. montenegrensis* | 23 | 24 | 6 | 28 | 2 | 30 |
| *R. nasutus* | 20 | 21 | 7 | 28 | 0 | 28 |
| *R. neglectus* | 23 | 25 | 1 | 26 | 0 | 26 |
| *R. neivai* | 30 | 32 | 5 | 36 | 1 | 37 |
| *R. pallescens* | 24 | 24 | 11 | 32 | 3 | 35 |
| *R. pictipes* | 28 | 31 | 13 | 41 | 3 | 44 |
| *R. prolixus* | 25 | 30 | 6 | 31 | 5 | 36 |
| *R. robustus* | 29 | 31 | 6 | 36 | 1 | 37 |
| IRs (before manual curation) |  |  |  |  |  |  |
| *R. brethesi* | 31 | 42 | 15 | 51 | 6 | 57 |
| *R. colombiensis* | 35 | 43 | 20 | 57 | 6 | 63 |
| *R. coreodes* | 32 | 47 | 14 | 51 | 10 | 61 |
| *R. domesticus* | 34 | 38 | 6 | 41 | 3 | 44 |
| *R. montenegrensis* | 26 | 31 | 10 | 40 | 1 | 41 |
| *R. milesi* | 32 | 41 | 13 | 46 | 8 | 54 |
| *R. nasutus* | 18 | 24 | 16 | 33 | 7 | 40 |
| *R. neglectus* | 33 | 48 | 13 | 56 | 5 | 61 |
| *R. neivai* | 36 | 47 | 4 | 44 | 7 | 51 |
| *R. pallescens* | 30 | 43 | 17 | 53 | 7 | 60 |
| *R. pictipes* | 30 | 36 | 22 | 57 | 1 | 58 |
| *R. prolixus* | 30 | 37 | 23 | 58 | 2 | 60 |
| *R. robustus* | 33 | 46 | 15 | 51 | 10 | 61 |

**Supplementary Table 3:** Matrix of presence or absence of the different ORs by ortholog groups found in the 13 assemblies (in red the domiciliary species)

|  | *R.*<br>*breth*<br>*esi* | *R.*<br>*colo*<br>*mbie*<br>*nsis* | *R.*<br>*coreo*<br>*des* | *R.*<br>*dome*<br>*sticus* | *R.*<br>*milesi* | *R.*<br>*mont*<br>*enegr*<br>*ensis* | *R.*<br>*nasut*<br>*us* | *R.*<br>*negle*<br>*ctus* | *R.*<br>*neiva*<br>*i* | *R.*<br>*palles*<br>*cens* | *R.*<br>*pictip*<br>*es* | *R.*<br>*prolix*<br>*us* | *R.*<br>*robust*<br>*us* | TOTAL |
| --- | --- | --- | --- | --- | --- | --- | --- | --- | --- | --- | --- | --- | --- | --- |
| OR1 | 1 | 1 | 1 | 1 | 1 |  | 1 | 1 | 1 | 1 | 1 | 1 |  | 8 |
| OR2 | 1 | 1 | 1 | 1 | 1 |  |  | 1 | 1 | 1 |  | 1 | 1 | 7 |
| OR3-5 | 1 | 1 | 1 |  | 2 | 1 | 1 | 1 | 2 |  | 1 | 3 | 3 | 14 |
| OR7 | 1 | 1 | 2 | 1 | 1 |  | 1 | 2 | 1 |  | 2 | 1 | 2 | 11 |
| OR8 |  | 1 |  |  | 1 |  |  | 1 |  |  |  | 1 | 1 | 4 |
| OR9 |  | 1 | 1 | 1 | 1 | 1 |  | 1 | 1 |  |  |  | 1 | 6 |
| OR12 | 1 | 1 |  | 1 | 1 | 1 |  | 1 | 1 | 1 |  |  | 1 | 7 |
| OR13 | 1 | 1 | 1 | 1 | 1 |  | 1 | 1 | 1 |  | 1 | 1 | 1 | 8 |
| OR14 | 1 | 1 | 1 | 1 | 1 | 1 | 1 | 1 | 1 | 1 | 1 |  | 1 | 9 |
| OR16-18 | 1 | 1 | 2 | 1 | 3 | 2 |  | 2 | 3 |  |  | 3 | 3 | 17 |
| OR19 | 1 | 1 | 1 | 1 | 1 | 1 |  | 1 | 1 |  | 1 | 1 | 1 | 8 |
| OR20 | 1 | 1 | 1 | 1 | 1 | 1 |  | 1 | 1 |  |  | 1 | 1 | 7 |
| OR21-23 | 1 | 2 | 1 | 2 | 3 | 2 | 1 | 3 | 3 | 1 | 2 | 2 | 2 | 21 |
| OR24 | 1 | 1 | 1 | 1 | 1 | 1 |  | 1 | 1 | 1 | 2 |  | 1 | 9 |
| OR25 | 1 | 1 |  |  | 1 | 1 | 1 | 1 | 1 |  | 1 | 1 | 1 | 8 |
| OR26 | 1 | 1 | 1 | 1 | 1 | 1 |  | 1 | 1 | 1 | 1 | 1 | 1 | 9 |
| OR27 | 1 | 1 | 1 | 1 | 1 |  |  |  | 1 |  |  | 1 | 1 | 5 |
| OR28-29 | 2 | 7 | 3 | 4 | 4 | 6 |  | 3 | 5 |  | 4 | 4 | 7 | 37 |
| OR30 | 1 | 1 | 1 | 1 | 1 | 1 | 1 | 1 | 1 | 1 | 1 |  | 1 | 9 |
| OR31 | 1 | 1 | 1 | 1 | 1 | 1 | 1 | 1 | 1 |  | 1 | 1 |  | 8 |
| OR32 | 1 | 1 | 1 | 1 | 1 | 1 | 1 |  | 1 | 1 | 1 | 1 | 1 | 9 |
| OR33 |  | 1 |  |  | 1 |  |  |  | 1 |  | 1 |  | 1 | 4 |
| OR34 | 1 | 1 | 1 | 1 | 1 | 1 | 1 | 1 | 1 | 1 | 1 | 1 |  | 9 |
| OR36-37 | 1 |  | 1 | 1 | 2 | 1 |  | 2 | 1 | 1 |  |  | 2 | 10 |
| OR38 |  |  | 1 | 1 | 1 | 1 |  | 1 |  |  | 1 | 1 |  | 6 |
| OR39 | 1 | 1 | 1 | 1 | 1 | 2 | 1 | 1 | 1 |  | 1 | 1 | 1 | 10 |
| OR40 | 1 | 1 | 1 | 1 | 1 |  | 1 | 1 | 1 |  | 1 | 1 | 1 | 8 |
| OR41 | 1 | 1 | 1 |  |  |  | 1 | 1 | 1 |  | 1 | 1 | 1 | 6 |
| OR42 | 1 | 1 |  | 1 | 1 |  |  | 1 | 1 |  | 1 | 1 | 1 | 7 |
| OR43 | 1 | 1 | 1 | 1 | 1 |  | 1 | 1 | 1 |  |  | 1 | 1 | 7 |
| OR44 | 1 | 1 | 1 | 1 | 1 | 1 |  | 1 | 1 | 1 | 1 | 1 | 1 | 9 |
| OR45 | 1 | 1 | 1 | 1 | 1 | 1 |  | 1 | 1 |  | 1 | 1 | 1 | 8 |
| OR46 |  | 1 | 1 | 1 | 1 |  |  | 1 | 1 |  | 1 |  | 1 | 6 |
| OR47 |  | 1 | 1 | 1 | 1 | 1 |  | 1 | 1 |  | 1 |  | 1 | 7 |
| OR48 | 1 | 1 | 1 | 1 | 1 | 1 |  | 1 | 2 |  | 2 |  | 2 | 10 |
| OR49 | 1 | 1 | 1 | 1 | 1 | 1 | 1 | 1 |  | 1 | 1 | 1 | 1 | 9 |
| OR50 | 1 | 1 | 1 | 1 | 1 | 1 | 1 | 1 | 1 |  | 1 | 1 | 1 | 9 |
| OR51 | 1 | 1 | 1 | 1 | 1 | 1 |  |  | 1 | 1 | 1 | 1 | 1 | 8 |
| OR52 | 1 | 1 | 1 | 1 | 1 | 1 |  | 1 | 1 |  | 1 | 2 |  | 8 |
| OR53 | 1 | 1 | 1 | 1 | 1 | 1 |  |  | 1 |  |  | 1 | 1 | 6 |
| OR54 | 1 | 2 | 2 | 2 | 3 | 2 | 1 | 3 | 3 |  | 2 | 1 | 1 | 18 |
| OR55 | 1 | 1 | 1 | 1 |  | 1 |  | 1 | 1 |  | 1 | 1 | 1 | 7 |
| OR56 | 1 | 1 |  | 1 |  | 1 | 1 |  | 1 |  | 1 |  |  | 5 |
| OR57 | 1 | 1 | 1 | 1 | 1 | 1 |  | 1 | 1 |  |  | 1 |  | 6 |
| OR58 & 61 | 2 | 1 |  |  |  | 2 |  | 1 |  |  | 2 | 1 | 2 | 8 |
| OR59-60 | 2 | 2 |  | 2 | 1 | 1 |  | 2 |  |  |  | 1 | 1 | 8 |
| OR62 | 1 | 1 |  | 1 |  | 1 | 1 |  | 1 |  | 1 | 1 | 1 | 7 |
| OR63-69 |  |  |  |  |  |  |  |  |  |  |  |  |  | 0 |
| OR70 |  | 1 | 1 | 1 | 1 | 1 | 1 | 1 | 1 |  |  | 1 | 1 | 8 |
| OR71-73 |  |  |  |  |  |  |  |  | 3 |  | 1 |  |  | 4 |
| OR74-78 |  |  |  |  | 1 |  | 1 |  |  |  |  |  |  | 2 |
| OR79 | 1 | 1 | 1 | 1 | 1 | 1 |  | 1 | 1 |  | 1 | 1 | 1 | 8 |
| OR80 |  |  | 1 |  |  |  |  |  |  |  |  |  |  | 0 |
| OR81 |  |  |  |  |  |  |  |  |  |  |  |  |  | 0 |
| OR82 |  | 1 |  |  |  |  |  |  |  |  |  |  |  | 0 |
| OR83 |  |  |  |  |  |  |  |  |  |  |  | 1 | 1 | 2 |
| OR84 |  |  | 1 |  | 2 |  | 2 | 1 | 1 |  | 1 | 1 | 1 | 9 |
| OR85 | 1 |  |  |  | 1 | 1 |  |  | 1 |  | 2 | 1 |  | 6 |
| OR86-87 | 3 | 1 |  | 1 | 1 | 1 |  |  | 1 |  | 2 | 2 | 2 | 10 |
| OR88-91 |  |  |  |  |  |  |  |  |  |  |  |  |  | 0 |
| OR92 |  |  | 1 | 2 | 1 |  | 1 | 1 | 1 |  |  | 1 | 1 | 8 |
| OR93 |  |  |  |  |  | 1 |  |  |  |  |  |  |  | 1 |
| OR94 |  |  |  |  |  |  |  |  |  |  |  |  |  | 0 |
| OR95 |  |  |  |  | 1 | 1 | 1 | 1 |  |  |  | 1 | 1 | 6 |
| OR96 |  | 1 | 1 | 1 |  |  |  |  | 1 | 1 | 1 |  | 1 | 5 |
| OR97 | 1 | 1 | 1 | 1 | 1 | 1 |  | 1 | 1 | 1 |  |  | 1 | 7 |
| OR98 | 1 | 1 | 1 |  | 1 | 1 |  | 1 | 1 |  |  |  | 1 | 5 |
| OR99 |  | 1 | 1 | 1 | 1 | 1 | 1 | 1 | 1 |  | 1 | 1 | 1 | 9 |
| OR100 |  |  |  |  |  |  |  |  |  |  |  |  |  | 0 |
| OR101 | 1 | 1 | 1 | 1 | 1 | 1 | 1 | 1 | 1 |  | 1 | 1 | 1 | 9 |
| OR102 | 1 | 1 | 1 | 1 | 1 | 1 |  | 1 | 1 | 1 | 1 | 1 | 1 | 9 |
| OR103 |  | 1 | 1 | 1 |  |  | 1 | 1 | 1 |  | 1 |  | 1 | 6 |
| OR104 |  |  | 1 |  | 1 | 1 |  | 1 | 1 |  |  |  |  | 4 |
| OR105 |  |  |  |  |  |  |  |  |  |  |  | 1 | 1 | 2 |
| OR106 | 1 | 1 | 1 | 1 | 1 |  | 1 | 1 | 1 |  | 1 | 1 | 1 | 8 |
| OR107 |  |  |  | 1 |  |  |  |  | 1 |  |  | 1 |  | 3 |
| OR108 | 1 | 1 | 1 | 1 | 1 | 1 |  | 1 | 1 |  | 1 | 1 |  | 7 |
| OR109 |  |  |  |  |  |  |  |  |  |  |  |  |  | 0 |
| OR110 |  |  | 1 | 1 |  | 1 |  | 1 | 1 | 1 | 1 | 1 |  | 7 |
| OR111 |  | 1 | 1 | 1 | 1 | 1 |  | 1 | 1 | 1 |  |  | 1 | 7 |
| OR112 | 2 |  |  | 2 | 1 |  |  | 2 |  |  | 1 | 1 | 1 | 8 |
| OR113 | 1 | 1 | 1 | 1 | 1 | 1 | 1 |  | 1 |  | 1 |  | 1 | 7 |
| OR114-115 | 1 | 1 | 2 | 2 | 2 | 2 |  |  | 2 | 1 | 1 | 1 | 2 | 13 |
| OR116 (NEW) |  | 1 |  | 1 | 1 | 1 |  |  | 1 | 1 |  |  | 1 | 6 |
| OR117 (NEW) |  | 1 | 1 | 1 | 1 | 1 |  | 1 |  | 1 | 1 | 1 | 1 | 8 |
| OR118 (NEW) |  | 1 | 1 | 1 | 1 | 1 | 1 | 1 | 1 | 1 | 1 | 1 |  | 9 |
| OR119 (NEW) | 1 | 1 |  |  |  |  |  |  |  |  | 1 |  |  | 1 |
| OR120 (NEW) |  | 1 | 1 |  | 1 | 1 | 1 | 2 |  |  | 1 | 1 | 1 | 8 |
| OR121 (NEW) | 1 | 1 | 2 |  | 1 |  |  |  | 2 |  |  | 1 |  | 4 |
| OR122 (NEW) | 4 | 1 | 1 |  | 1 | 2 |  | 1 |  |  | 4 |  | 2 | 10 |
| ORco | 1 | 1 | 1 | 1 | 1 | 1 | 1 | 1 | 1 | 1 |  | 1 | 1 | 9 |
| TOTAL | 66 | 78 | 70 | 72 | 82 | 70 | 34 | 74 | 84 | 24 | 70 | 69 | 83 | 662 |

**Supplementary Table 4:**
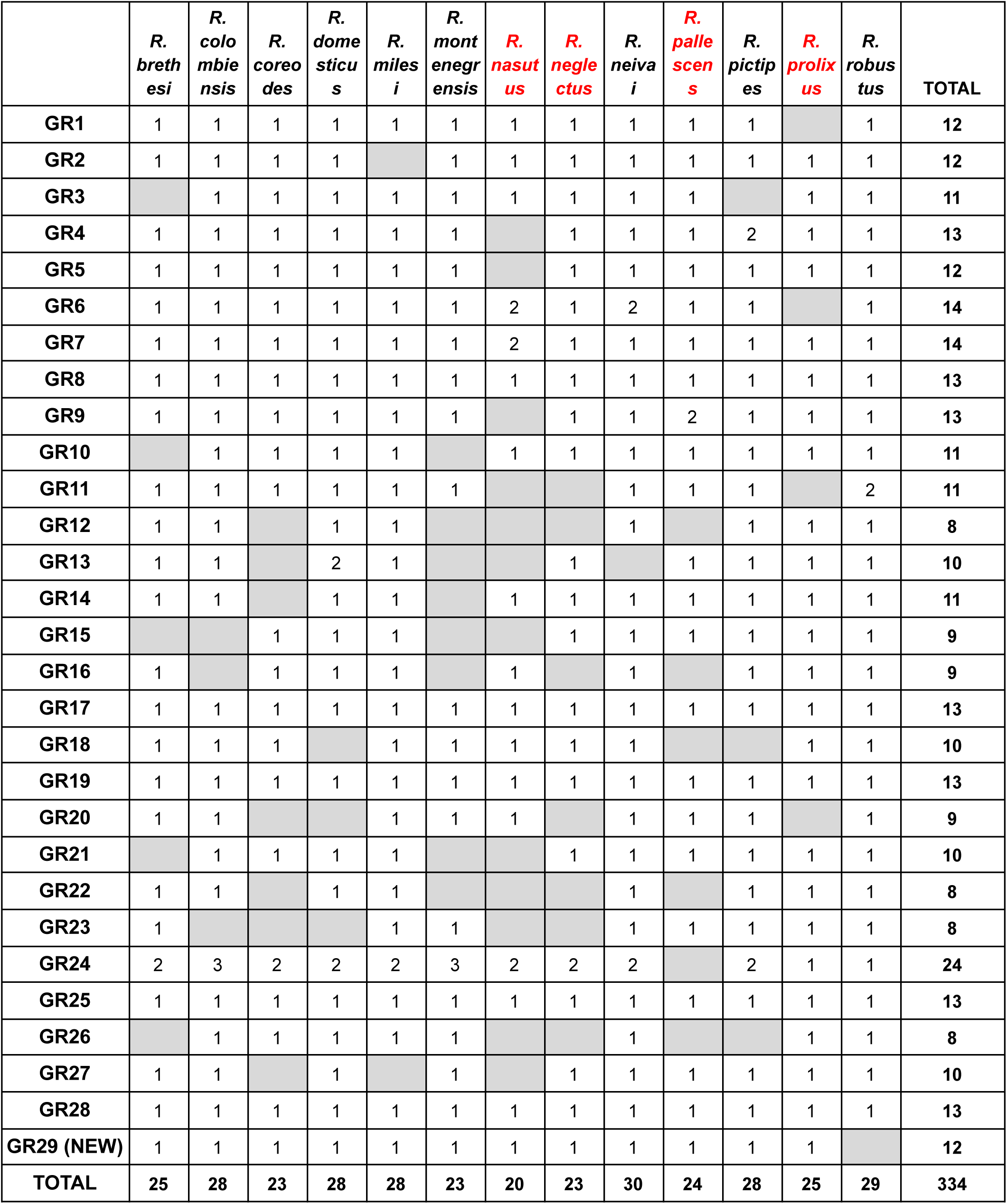
Matrix of presence or absence of the different GRs by ortholog groups found in the 13 assemblies (in red the domiciliary species).

|  | *R. brethesi* | *R. colombiensis* | *R. coreodes* | *R. domesticus* | *R. milesi* | *R. montenegrensis* | *R. nasutus* | *R. neglectus* | *R. neivai* | *R. pallens* | *R. pictipes* | *R. prolixus* | *R. robustus* | TOTAL |
| --- | --- | --- | --- | --- | --- | --- | --- | --- | --- | --- | --- | --- | --- | --- |
| GR1 | 1 | 1 | 1 | 1 | 1 | 1 | 1 | 1 | 1 | 1 | 1 |  | 1 | 12 |
| GR2 | 1 | 1 | 1 | 1 |  | 1 | 1 | 1 | 1 | 1 | 1 | 1 | 1 | 12 |
| GR3 |  | 1 | 1 | 1 | 1 | 1 | 1 | 1 | 1 | 1 |  | 1 | 1 | 11 |
| GR4 | 1 | 1 | 1 | 1 | 1 | 1 |  | 1 | 1 | 1 | 2 | 1 | 1 | 13 |
| GR5 | 1 | 1 | 1 | 1 | 1 | 1 |  | 1 | 1 | 1 | 1 | 1 | 1 | 12 |
| GR6 | 1 | 1 | 1 | 1 | 1 | 1 | 2 | 1 | 2 | 1 | 1 |  | 1 | 14 |
| GR7 | 1 | 1 | 1 | 1 | 1 | 1 | 2 | 1 | 1 | 1 | 1 | 1 | 1 | 14 |
| GR8 | 1 | 1 | 1 | 1 | 1 | 1 | 1 | 1 | 1 | 1 | 1 | 1 | 1 | 13 |
| GR9 | 1 | 1 | 1 | 1 | 1 | 1 |  | 1 | 1 | 2 | 1 | 1 | 1 | 13 |
| GR10 |  | 1 | 1 | 1 | 1 |  | 1 | 1 | 1 | 1 | 1 | 1 | 1 | 11 |
| GR11 | 1 | 1 | 1 | 1 | 1 | 1 |  |  | 1 | 1 | 1 |  | 2 | 11 |
| GR12 | 1 | 1 |  | 1 | 1 |  |  |  | 1 |  | 1 | 1 | 1 | 8 |
| GR13 | 1 | 1 |  | 2 | 1 |  |  | 1 |  | 1 | 1 | 1 | 1 | 10 |
| GR14 | 1 | 1 |  | 1 | 1 |  | 1 | 1 | 1 | 1 | 1 | 1 | 1 | 11 |
| GR15 |  |  | 1 | 1 | 1 |  |  | 1 | 1 | 1 | 1 | 1 | 1 | 9 |
| GR16 | 1 |  | 1 | 1 | 1 |  | 1 |  | 1 |  | 1 | 1 | 1 | 9 |
| GR17 | 1 | 1 | 1 | 1 | 1 | 1 | 1 | 1 | 1 | 1 | 1 | 1 | 1 | 13 |
| GR18 | 1 | 1 | 1 |  | 1 | 1 | 1 | 1 | 1 |  |  | 1 | 1 | 10 |
| GR19 | 1 | 1 | 1 | 1 | 1 | 1 | 1 | 1 | 1 | 1 | 1 | 1 | 1 | 13 |
| GR20 | 1 | 1 |  |  | 1 | 1 | 1 |  | 1 | 1 | 1 |  | 1 | 9 |
| GR21 |  | 1 | 1 | 1 | 1 |  |  | 1 | 1 | 1 | 1 | 1 | 1 | 10 |
| GR22 | 1 | 1 |  | 1 | 1 |  |  |  | 1 |  | 1 | 1 | 1 | 8 |
| GR23 | 1 |  |  |  | 1 | 1 |  |  | 1 | 1 | 1 | 1 | 1 | 8 |
| GR24 | 2 | 3 | 2 | 2 | 2 | 3 | 2 | 2 | 2 |  | 2 | 1 | 1 | 24 |
| GR25 | 1 | 1 | 1 | 1 | 1 | 1 | 1 | 1 | 1 | 1 | 1 | 1 | 1 | 13 |
| GR26 |  | 1 | 1 | 1 | 1 | 1 |  |  | 1 |  |  | 1 | 1 | 8 |
| GR27 | 1 | 1 |  | 1 |  | 1 |  | 1 | 1 | 1 | 1 | 1 | 1 | 10 |
| GR28 | 1 | 1 | 1 | 1 | 1 | 1 | 1 | 1 | 1 | 1 | 1 | 1 | 1 | 13 |
| GR29 (NEW) | 1 | 1 | 1 | 1 | 1 | 1 | 1 | 1 | 1 | 1 | 1 | 1 |  | 12 |
| TOTAL | 25 | 28 | 23 | 28 | 28 | 23 | 20 | 23 | 30 | 24 | 28 | 25 | 29 | 334 |

**Supplementary Table 5:**
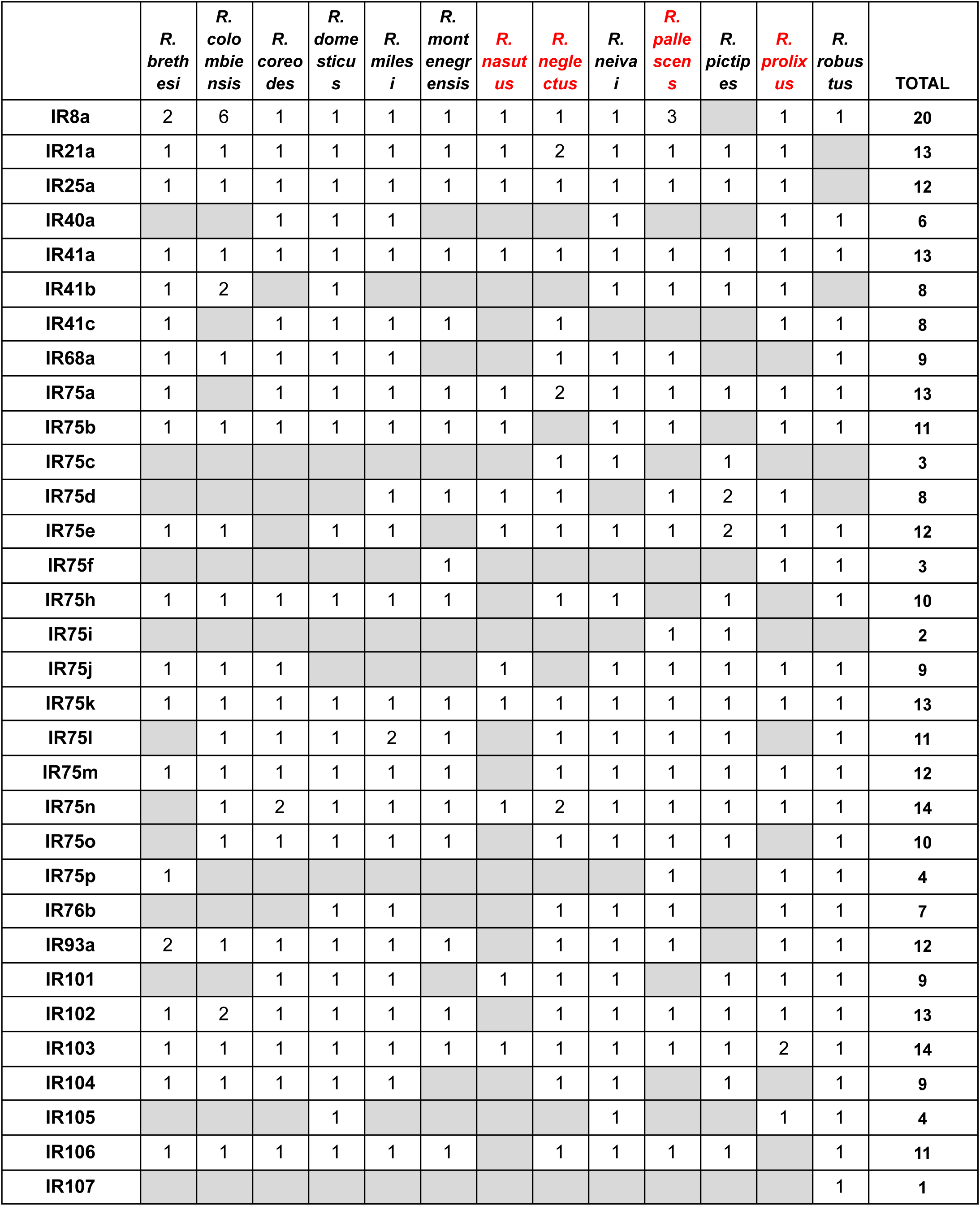
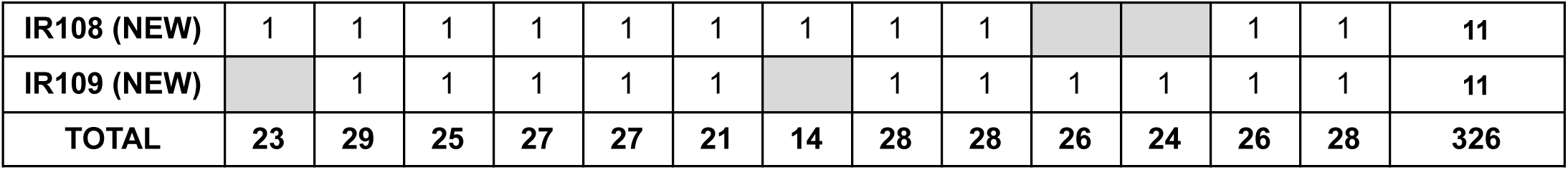
Matrix of presence or absence of the different IRs by ortholog groups found in the 13 assemblies (in red the domiciliary species).

**Supplementary Table 6:**
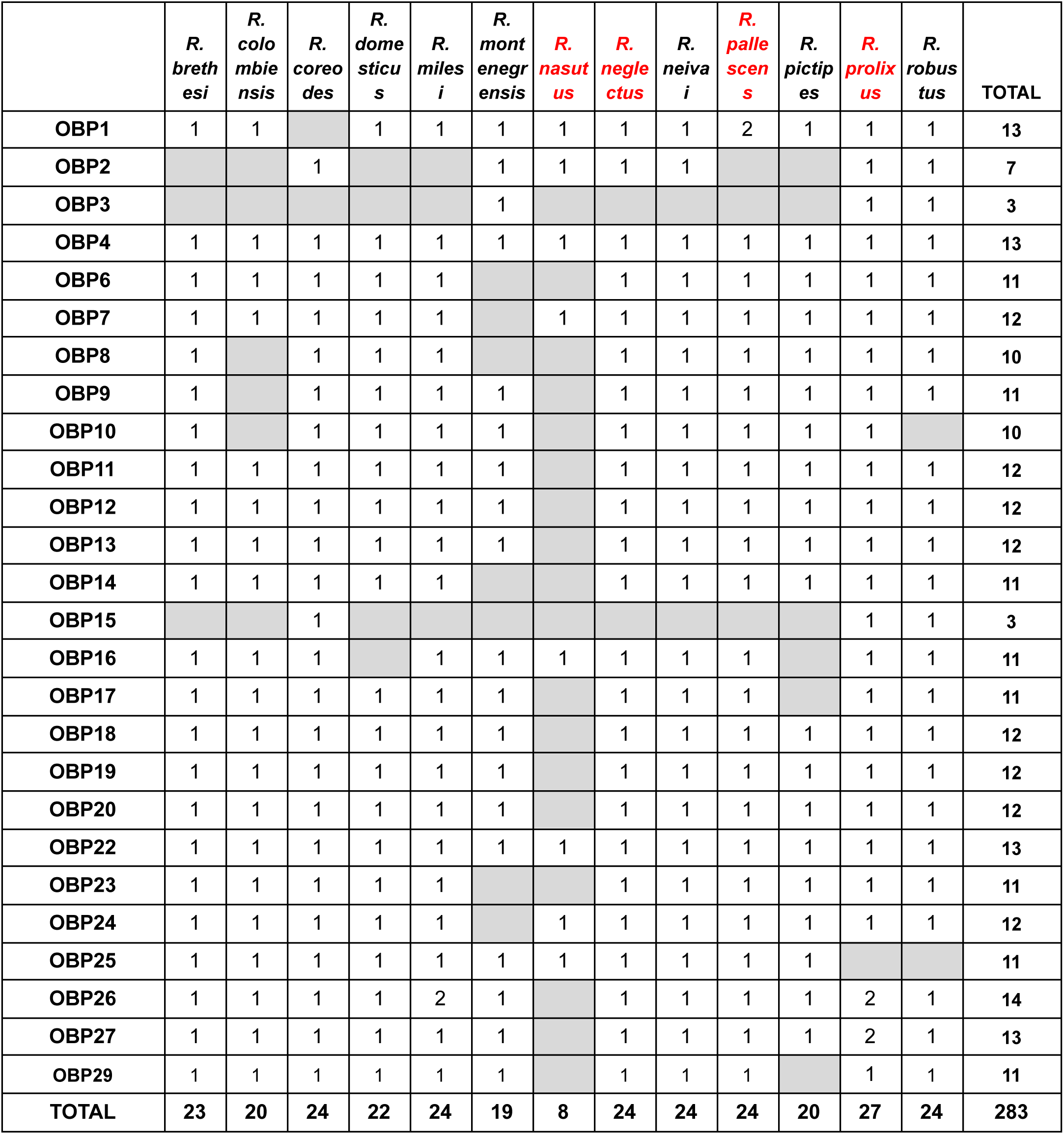
Matrix of presence or absence of the different OBPs by ortholog groups found in the 13 assemblies (in red the domiciliary species).

**Supplementary Table 7:** Matrix of presence or absence of the different CSPs by ortholog groups found in the 13 assemblies (in red the domiciliary species).

|  | *R.*<br>*breth*<br>*esi* | *R.*<br>*colo*<br>*mbie*<br>*nsis* | *R.*<br>*coreo*<br>*des* | *R.*<br>*dome*<br>*sticu*<br>*s* | *R.*<br>*miles*<br>*i* | *R.*<br>*mont*<br>*enegr*<br>*ensis* | *R.*<br>*nasut*<br>*us* | *R.*<br>*negle*<br>*ctus* | *R.*<br>*neiva*<br>*i* | *R.*<br>*palle*<br>*scen*<br>*s* | *R.*<br>*pictip*<br>*es* | *R.*<br>*prolix*<br>*us* | *R.*<br>*robus*<br>*tus* | TOTAL |
| --- | --- | --- | --- | --- | --- | --- | --- | --- | --- | --- | --- | --- | --- | --- |
| CSP1 | 1 |  | 1 | 1 | 1 |  | 1 | 1 | 1 | 1 | 1 | 1 | 1 | 11 |
| CSP2 | 1 | 1 | 1 | 1 | 1 |  | 1 | 1 | 1 | 1 | 1 | 1 | 1 | 12 |
| CSP3 | 1 | 1 | 1 | 1 | 1 | 1 | 1 | 1 | 1 | 1 | 1 | 1 | 1 | 13 |
| CSP4 | 1 | 1 | 1 | 1 | 1 |  | 1 | 1 | 1 | 1 | 1 | 1 | 1 | 12 |
| CSP5 | 1 | 1 | 1 | 1 | 1 | 1 | 1 | 1 | 1 | 1 | 1 | 1 | 1 | 13 |
| CSP6 |  | 1 | 1 | 1 | 1 | 1 | 1 | 1 | 1 | 1 | 1 | 1 | 1 | 12 |
| CSP7 | 1 | 1 | 1 | 1 | 1 | 1 | 1 | 1 | 1 | 1 | 1 | 1 | 1 | 13 |
| CSP8 |  | 1 | 1 | 1 | 1 |  |  |  | 1 |  |  |  | 1 | 6 |
| CSP9 |  | 1 | 1 |  | 1 |  |  | 1 |  |  |  | 1 | 1 | 6 |
| CSP10 | 1 | 1 | 1 | 1 | 1 |  |  | 1 | 1 | 1 | 1 | 1 | 1 | 11 |
| CSP11 | 1 | 1 | 1 | 1 | 1 |  | 1 | 1 | 1 | 1 |  | 1 | 1 | 11 |
| CSP12 |  | 1 | 1 | 1 | 1 | 1 |  | 1 | 1 |  | 1 | 1 |  | 9 |
| CSP13 |  | 1 | 1 | 1 | 1 | 1 |  | 1 | 1 | 1 | 1 | 1 | 1 | 11 |
| CSP14-15 | 1 | 1 | 1 | 1 | 1 |  | 1 | 1 | 1 |  |  |  | 1 | 9 |
| CSP16 |  |  |  | 1 | 1 |  |  |  | 1 |  |  |  | 1 | 4 |
| CSP17 | 1 | 1 | 1 | 1 | 1 | 1 | 1 | 1 | 1 | 1 | 1 | 1 | 1 | 13 |
| CSP18 | 1 | 1 | 1 | 1 | 1 | 1 | 1 | 1 | 1 | 1 | 1 | 1 | 1 | 13 |
| CSP19 | 1 | 1 | 1 | 1 | 1 | 1 | 1 | 1 | 1 | 1 |  | 1 | 1 | 12 |
| CSP20 (NEW) | 1 | 1 | 1 | 1 | 1 | 1 | 1 | 1 | 1 | 1 | 1 | 1 | 1 | 13 |
| TOTAL | 13 | 17 | 18 | 18 | 19 | 10 | 13 | 17 | 18 | 14 | 13 | 16 | 18 | 204 |

**Supplementary Table 8:**
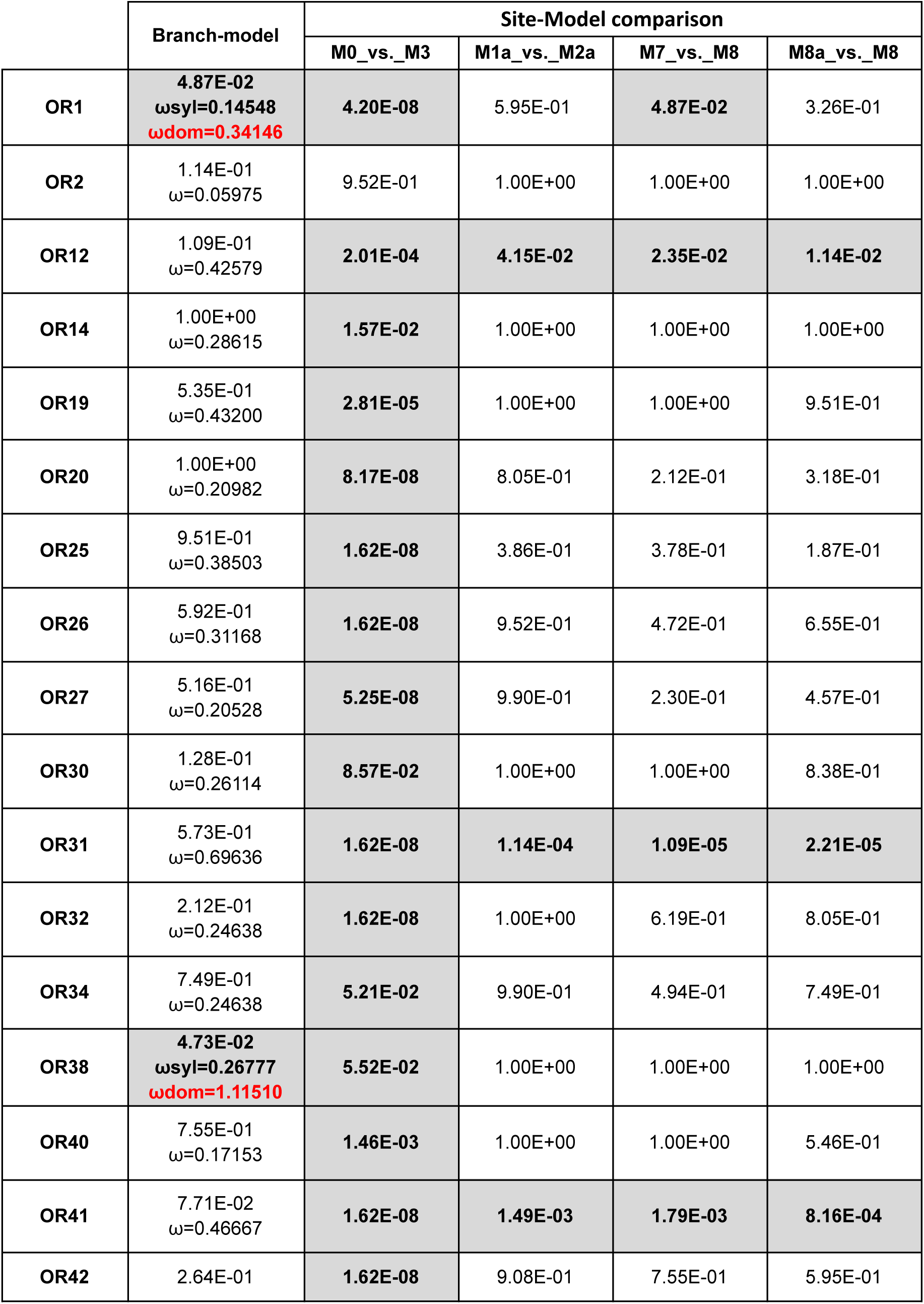
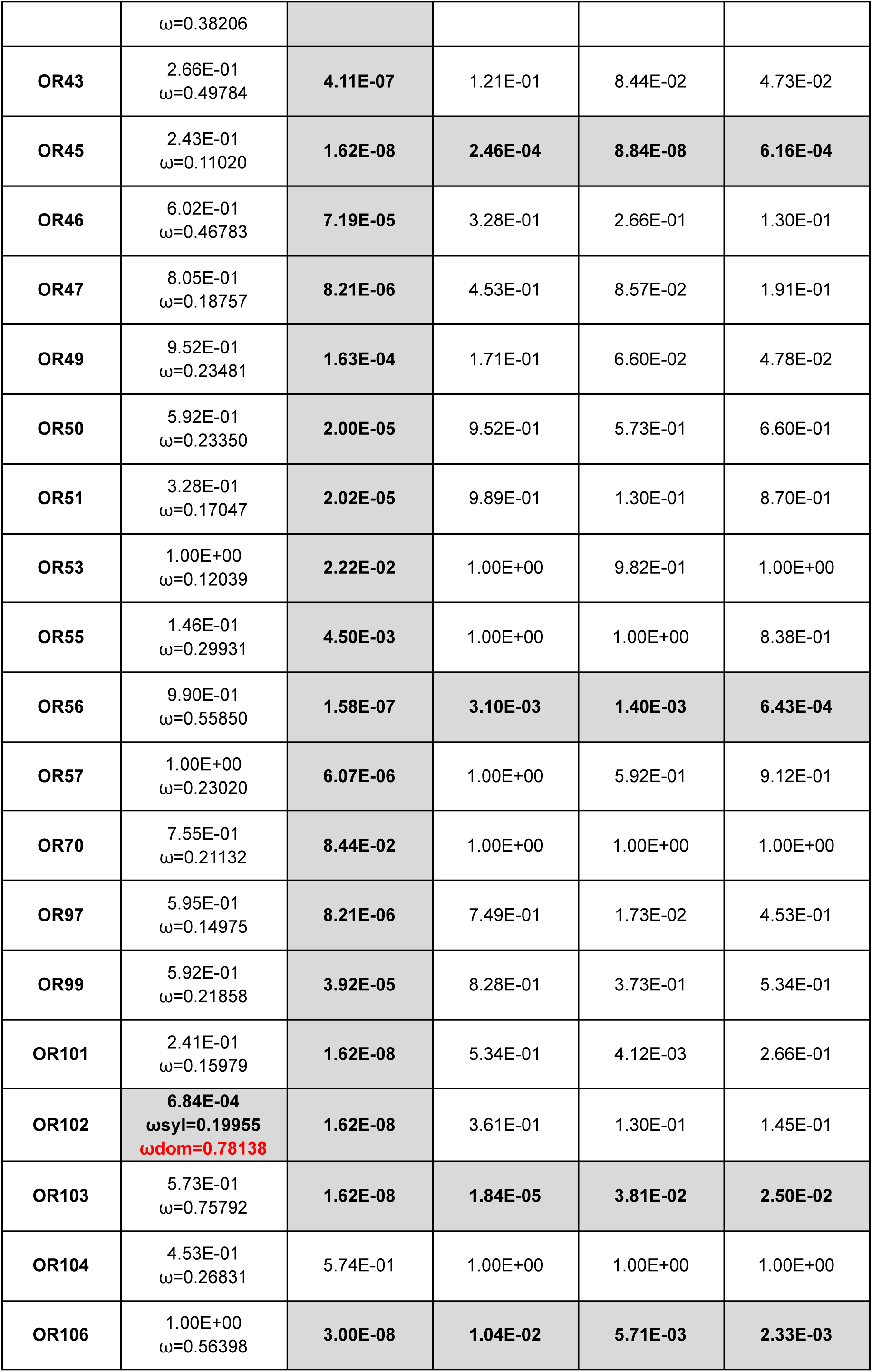
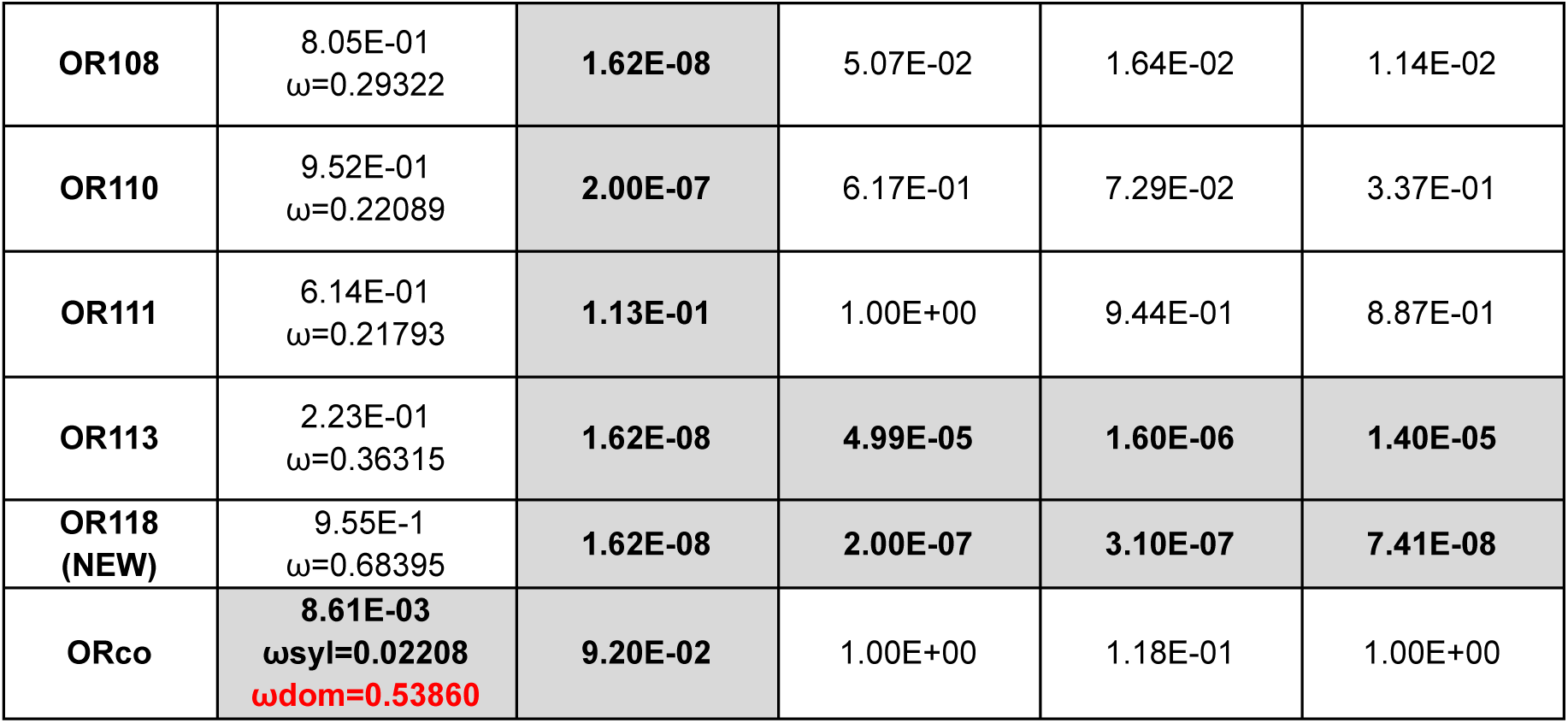
CodeML results for ORs. For each OR and each test, the adjusted p-value (according to Benjamini and Hochberg) is indicated. For significant branch model tests, dN/dS values are provided for sylvatic and domiciliary species separately. For non-significant tests, the global dN/dS value across the entire tree is reported.

**Supplementary Table 9:** CodeML results for GRs. For each GR and each test, the adjusted p-value (according to Benjamini and Hochberg) is indicated. For significant branch model tests, dN/dS values are provided for sylvatic and domiciliary species separately. For non-significant tests, the global dN/dS value across the entire tree is reported. Independent analyses were performed for GR6 which contains two duplicated copies in *R. neivai* and for the two genes GR11 and GR13, each of which contains two duplicated copies in *R. robustus*

|  | Branch-Model | Site-Model comparison |  |  |  |
| --- | --- | --- | --- | --- | --- |
|  |  | M0_vs._M3 | M1a_vs._M2a | M7_vs._M8 | M8a_vs._M8 |
| GR1 | 1.47E-01<br>$\omega=0.09125$ | 1.00E+00 | 1.00E+00 | 1.00E+00 | 8.98E-01 |
| GR2 | 2.78E-01<br>$\omega=0.09409$ | 2.18E-01 | 1.00E+00 | 1.00E+00 | 7.43E-01 |
| GR3 | 1.04E-01<br>$\omega=0.25523$ | 1.00E+00 | 1.00E+00 | 1.00E+00 | 1.00E+00 |
| GR4 | 9.57E-01<br>$\omega=0.14568$ | 3.39E-02 | 1.00E+00 | 6.09E-01 | 8.26E-01 |
| GR5 | 4.89E-01<br>$\omega=0.55392$ | 2.30E-04 | 1.88E-01 | 1.04E-01 | 8.38E-02 |
| GR6 (*R. neivai* dup1) | 1.36E-01<br>$\omega=0.55392$ | 2.53E-06 | 3.09E-01 | 2.78E-01 | 1.40E-01 |
| GR6 (*R. neivai* dup2) | 1.67E-01<br>$\omega=0.46374$ | 3.91E-06 | 2.17E-01 | 1.66E-01 | 9.92E-02 |
| GR7 | 1.00E+00<br>$\omega=0.26108$ | 3.00E-08 | 2.90E-01 | 3.95E-02 | 1.19E-01 |
| GR8 | 3.06E-01<br>$\omega=0.17440$ | 2.84E-06 | 1.00E+00 | 7.29E-01 | 1.00E+00 |
| GR9 | 3.09E-01<br>$\omega=0.17705$ | 2.84E-06 | 1.00E+00 | 7.29E-01 | 1.00E+00 |
| GR10 | 1.00E+00<br>$\omega=0.31022$ | 6.06E-05 | 2.96E-01 | 1.21E-01 | 8.38E-02 |
| GR11 (*R. robustus* dup1) | 6.49E-01<br>$\omega=0.58604$ | 3.00E-08 | 2.30E-04 | 7.25E-05 | 6.76E-05 |
| GR11 (*R. robustus* dup2) | 5.40E-01<br>$\omega=0.54703$ | 3.00E-08 | 6.90E-04 | 2.30E-04 | 1.49E-04 |
| GR12 | 1.46E-01<br>$\omega=0.49059$ | 3.50E-02 | 9.97E-01 | 7.29E-01 | 7.29E-01 |
| GR13 (*R. domesticus* dup1) | 3.09E-01<br>$\omega=0.62374$ | 1.15E-05 | 3.28E-01 | 2.17E-01 | 1.60E-01 |
| **GR13 (*R. domesticus* dup2)** | 2.84E-01<br>$\omega=0.64962$ | **4.00E-07** | 1.39E-01 | 9.31E-02 | 5.64E-02 |
| **GR14** | 2.17E-01<br>$\omega=0.64962$ | **3.61E-06** | 6.61E-01 | 4.92E-01 | 3.02E-01 |
| **GR15** | 1.40E-01<br>$\omega=0.06724$ | **3.95E-02** | 1.00E+00 | 4.92E-01 | 1.00E+00 |
| **GR16** | 4.17E-01<br>$\omega=0.18838$ | **3.00E-08** | 6.54E-01 | 1.12E-01 | 4.92E-01 |
| **GR17** | 7.31E-01<br>$\omega=0.10735$ | **7.45E-05** | 1.00E+00 | 1.00E+00 | 5.04E-01 |
| **GR18** | 8.61E-01<br>$\omega=0.14446$ | **2.03E-01** | 1.00E+00 | 1.00E+00 | 9.99E-01 |
| **GR19** | 1.00E+00<br>$\omega=0.37532$ | **8.19E-02** | 1.00E+00 | 9.99E-01 | 8.61E-01 |
| **GR20** | 2.60E-01<br>$\omega=0.39688$ | **8.53E-03** | 1.00E+00 | 1.00E+00 | 7.57E-01 |
| **GR21** | 7.33E-01<br>$\omega=0.29678$ | **1.19E-04** | 2.60E-01 | 1.22E-01 | 7.36E-02 |
| **GR22** | 1.00E+00<br>$\omega=0.10999$ | **1.77E-02** | 1.00E+00 | 5.84E-01 | 7.61E-01 |
| **GR25** | 9.31E-02<br>$\omega=0.26563$ | **2.30E-04** | 7.30E-01 | 4.51E-01 | 4.34E-01 |
| **GR26** | 1.00E+00<br>$\omega=0.30665$ | **7.73E-01** | 7.29E-01 | 5.89E-01 | 4.29E-01 |
| **GR27** | 8.76E-01<br>$\omega=0.21591$ | **2.99E-05** | 8.26E-01 | 3.73E-01 | 4.70E-01 |
| **GR28** | 7.35E-01<br>$\omega=0.30947$ | **3.23E-05** | 5.06E-01 | 3.80E-01 | 2.17E-01 |
| **GR29 (NEW)** | 9.99E-01<br>$\omega=0.47073$ | **3.00E-08** | **1.24E-05** | **1.15E-05** | **2.84E-06** |

**Supplementary Table 10:** CodeML results for IRs. For each IR and each test, the adjusted p-value (according to Benjamini and Hochberg) is indicated. For significant branch model tests, dN/dS values are provided for sylvatic and domiciliary species separately. For non-significant tests, the global dN/dS value across the entire tree is reported. Independent analyses were performed for IR75e which contains two copies in *R. pictipes* and for IR75n, each of which contains two duplicated copies in *R. coreodes*

|  | Branch-model | Site-Model comparison |  |  |  |
| --- | --- | --- | --- | --- | --- |
|  |  | M0_vs._M3 | M1a_vs._M2a | M7_vs._M8 | M8a_vs._M8 |
| IR21a | 1.28E-01<br>$\omega=0.34230$ | 1.04E-08 | 1.04E-08 | 1.04E-08 | 1.04E-08 |
| IR25a | 1.00E+00<br>$\omega=0.03296$ | 1.60E-04 | 9.44E-01 | 3.82E-03 | 6.47E-01 |
| IR40a | 1.00E+00<br>$\omega=0.03346$ | 1.00E+00 | 1.00E+00 | 9.92E-01 | 1.00E+00 |
| IR41a | 1.00E+00<br>$\omega=0.33669$ | 3.57E-01 | 1.00E+00 | 1.00E+00 | 1.00E+00 |
| IR41b | 5.80E-01<br>$\omega=0.22520$ | 1.69E-02 | 9.53E-01 | 6.53E-01 | 6.53E-01 |
| IR41c | 5.40E-01<br>$\omega=0.32581$ | 1.17E-02 | 1.00E+00 | 1.00E+00 | 1.00E+00 |
| IR68a | 3.43E-01<br>$\omega=0.07841$ | 1.35E-04 | 9.55E-01 | 4.15E-02 | 6.53E-01 |
| IR75a | 2.82E-02<br>$\omega_{syl}=0.18714$<br>$\omega_{dom}=0.00010$ | 9.72E-06 | 6.49E-01 | 1.77E-01 | 2.55E-01 |
| IR75b | 5.29E-01<br>$\omega=0.26602$ | 1.00E+00 | 1.00E+00 | 1.00E+00 | 1.00E+00 |
| IR75e (*R. pictipes* dup 1) | 7.51E-01<br>$\omega=0.15499$ | 1.04E-08 | 5.40E-01 | 6.22E-04 | 2.55E-01 |
| IR75e (*R. pictipes* dup 2) | 7.51E-01<br>$\omega=0.15670$ | 1.04E-08 | 5.40E-01 | 8.27E-04 | 2.55E-01 |
| IR75h | 1.00E+00<br>$\omega=0.01885$ | 1.00E+00 | 1.00E+00 | 1.00E+00 | 1.00E+00 |
| IR75j | 9.44E-01<br>$\omega=0.22328$ | 1.04E-08 | 8.55E-01 | 3.68E-02 | 5.40E-01 |
| IR75k | 1.00E+00<br>$\omega=0.20513$ | 1.04E-08 | 1.00E+00 | 1.00E+00 | 8.55E-01 |
| IR75l | 8.21E-01<br>$\omega=0.20850$ | 1.04E-08 | 6.20E-01 | 4.47E-02 | 4.44E-01 |
| IR75m | 7.51E-01<br>$\omega=0.32076$ | **1.04E-08** | 5.66E-02 | **9.45E-03** | **1.22E-02** |
| IR75n (*R. coreodes* dup1) | 9.44E-01<br>$\omega=0.15402$ | **4.73E-05** | 8.72E-01 | 1.06E-01 | 5.42E-01 |
| IR75n (*R. coreodes* dup2) | 9.44E-01<br>$\omega=0.15754$ | **7.42E-05** | 8.72E-01 | 1.09E-01 | 5.40E-01 |
| IR75o | 7.51E-01<br>$\omega=0.35891$ | **1.04E-08** | **4.59E-02** | **4.15E-02** | **2.35E-02** |
| IR75p | 6.38E-01<br>$\omega=0.36165$ | 8.56E-01 | 1.00E+00 | 1.00E+00 | 1.00E+00 |
| IR93a | 5.40E-01<br>$\omega=0.08074$ | **1.60E-04** | 1.00E+00 | 7.51E-01 | 3.16E-01 |
| IR101 | **3.63E-02**<br>$\omega_{syl}=0.16702$<br>$\omega_{dom}=2.24513$ | **1.20E-03** | 1.00E+00 | 8.72E-01 | 1.00E+00 |
| IR102 | 9.44E-01<br>$\omega=0.23489$ | **1.04E-08** | **3.68E-02** | **2.33E-04** | **7.76E-03** |
| IR103 | 1.00E+00<br>$\omega=0.20635$ | 3.79E-01 | 1.00E+00 | 1.00E+00 | 8.98E-01 |
| IR104 | 8.89E-01<br>$\omega=0.24189$ | **1.04E-08** | 1.00E+00 | 5.00E-01 | 9.56E-01 |
| IR105 | 1.00E+00<br>$\omega=0.29938$ | **1.36E-05** | **4.23E-02** | **2.82E-02** | **1.39E-02** |
| IR106 | 4.02E-01<br>$\omega=0.14926$ | **1.04E-08** | 5.40E-01 | **2.42E-03** | 2.55E-01 |
| IR108 (NEW) | 1.00E+00<br>$\omega=0.29899$ | 3.43E-01 | 1.00E+00 | 1.00E+00 | 1.00E+00 |
| IR109 (NEW) | 4.02E-01<br>$\omega=0.14754$ | **1.22E-04** | 2.78E-01 | 2.82E-02 | 1.20E-01 |

